# Allele-selective knockdown of mutant DNAJB6 improves molecular features in a primary mouse myotube model of LGMDD1

**DOI:** 10.64898/2026.11.06.764697

**Authors:** Aliasgar Vohra, Meade Haller, May M Paing, Alina Bao Lan Dam, Jil A Daw, Rocio Bengoechea, Sara K Pittman, Ting-Yu Wang, Carla M Yuede, Timothy M Miller, Heather L True, Scott Q Harper, Tsui-Fen Chou, Conrad C Weihl, Andrew R Findlay

## Abstract

Dominant variants in DNAJB6 cause limb-girdle muscular dystrophy D1 (LGMDD1), a progressive myopathy associated with disrupted protein homeostasis. Because complete loss of DNAJB6 is embryonic lethal in mice, we evaluated allele-selective knockdown as a strategy to reduce mutant DNAJB6 while preserving wild-type protein. We generated a knock-in mouse carrying an N-terminal 3×FLAG tag on the wild-type Dnajb6 allele, allowing endogenous wild-type and F90I DNAJB6, the murine equivalent of human F89I, to be measured within the same sample. Flag-LGMDD1 mice recapitulated the functional, myopathological, and biochemical features seen in other LGMDD1 models and in human disease. F90I DNAJB6 was more abundant and degraded more slowly than wild-type DNAJB6, and LGMDD1 muscle and primary myotubes showed altered protein handling at baseline and following proteostatic stress. Comparative proteomic analyses identified DNAJA4 as a candidate marker responsive to pathogenic DNAJB6 activity. A two-stage mismatch screen identified a lead candidate siRNA with 264-fold selectivity for F90I DNAJB6. In primary Flag-LGMDD1 myotubes, this siRNA selectively reduced F90I DNAJB6 and shifted several baseline and stress-dependent molecular abnormalities involving p62, HSP70, DNAJA4, and markers associated with unfolded protein response signaling toward control values. These findings establish cellular proof of concept for allele-selective knockdown in LGMDD1.

## INTRODUCTION

DNAJB6 is an HSP40 co-chaperone that works with HSP70 to maintain protein homeostasis. DNAJB6 binds misfolded client proteins and, through its J domain, stimulates HSP70 ATPase activity, promoting client transfer and either refolding or degradation. DNAJB6 is expressed as two major isoforms: the longer, predominantly nuclear DNAJB6a and the shorter, cytoplasmic DNAJB6b, which localizes to myofibrillar structures in skeletal muscle.^1,2^ Dominant pathogenic variants in DNAJB6 cause limb-girdle muscular dystrophy type D1 (LGMDD1), a progressive myopathy characterized by proximal weakness and myofibrillar pathology, including protein aggregates and rimmed vacuoles.^3–5^ Although DNAJB6 is ubiquitously expressed, the clinical phenotype is largely restricted to skeletal muscle. LGMDD1-associated mutations impair the interaction between the J and G/F domains of DNAJB6, relieving structural autoinhibition and allowing the J domain to interact with and stimulate HSP70 ATPase activity in the absence of client protein.^6^ Most pathogenic variants lie within the G/F domain, and a smaller number within the J domain disrupt the same interaction.^5,6^ This inappropriate activation is thought to sequester HSP70 and reduce its availability for other protein quality-control functions, creating a functional HSP70 deficiency.^6,7^ Abrogation of this unproductive interaction between LGMDD1 associated mutant DNAJB6 and HSP70 using small molecules or genetic manipulation is effective in rescuing phenotypes in LGMDD1 models.^8^

Selective suppression of a pathogenic allele is a potential therapeutic strategy for dominantly inherited disorders. In LGMDD1, however, nonselective suppression of DNAJB6 may carry substantial risk as complete loss of Dnajb6 is embryonic lethal in mice, and loss of dnajb6 in zebrafish skeletal muscle has been associated with mitochondrial abnormalities.^9,10^ By contrast, Dnajb6 haploinsufficiency is largely tolerated in mice, which do not develop skeletal muscle disease, although heterozygous mice show a mild, age-associated sinoatrial node phenotype in the absence of structural or functional cardiac remodeling.^11^ These findings suggest that selective suppression of mutant DNAJB6, while preserving wild-type expression, may provide a therapeutic window for LGMDD1. Here, we generated a FLAG-tagged knock-in mouse model that permits direct resolution of endogenous wild-type and mutant DNAJB6. Using this model, we characterized proteostasis abnormalities in skeletal muscle and primary myotubes, used proteomics to look for markers of pathogenic DNAJB6 activity, developed a systematic siRNA screening strategy for mutant-selective knockdown, and identified siRNAs with greater than 250-fold selectivity for mutant DNAJB6. We then evaluated whether selective reduction of mutant DNAJB6 corrects molecular features of LGMDD1 following proteostatic stress.

## RESULTS

### Flag-LGMDD1 mice permit allelic resolution of DNAJB6 and recapitulate features of LGMDD1

To directly measure endogenous wild-type and mutant DNAJB6 within the same sample, we used CRISPR/Cas9 to insert an N-terminal 3×FLAG tag into the wild-type *Dnajb6* allele (Fig. 1A,B). *Dnajb6* ^WT/WT-Flag^ mice were crossed with our previously characterized *Dnajb6* ^F90I/WT^ knock-in mice, which carry the murine equivalent of the human F89I mutation. In the resulting *Dnajb6*^F90I/WT-Flag^ mice, hereafter referred to as Flag-LGMDD1 mice, FLAG-tagged wild-type and untagged F90I DNAJB6 could be resolved by immunoblotting for both DNAJB6 isoforms (Fig. 1B). To assess whether the tag itself alters DNAJB6 abundance, we compared tagged and untagged species within myotube lysates from *Dnajb6*^WT/WT-Flag^ *and Dnajb6*^F90I/WT-Flag^ mice (Fig. S1). In *Dnajb6*^WT/WT-Flag^ lysates, where both alleles encode wild-type protein, tagged and untagged DNAJB6 appeared similar in abundance for both isoforms. In *Dnajb6*^F90I/WT-Flag^ lysates, untagged F90I DNAJB6 appeared more abundant than FLAG-WT DNAJB6 for both isoforms. This qualitative comparison provides no indication that FLAG tagging substantially alters DNAJB6 abundance and suggests that the increased DNAJB6 in Flag-LGMDD1 mice reflects the mutant allele.

**Figure 1.**
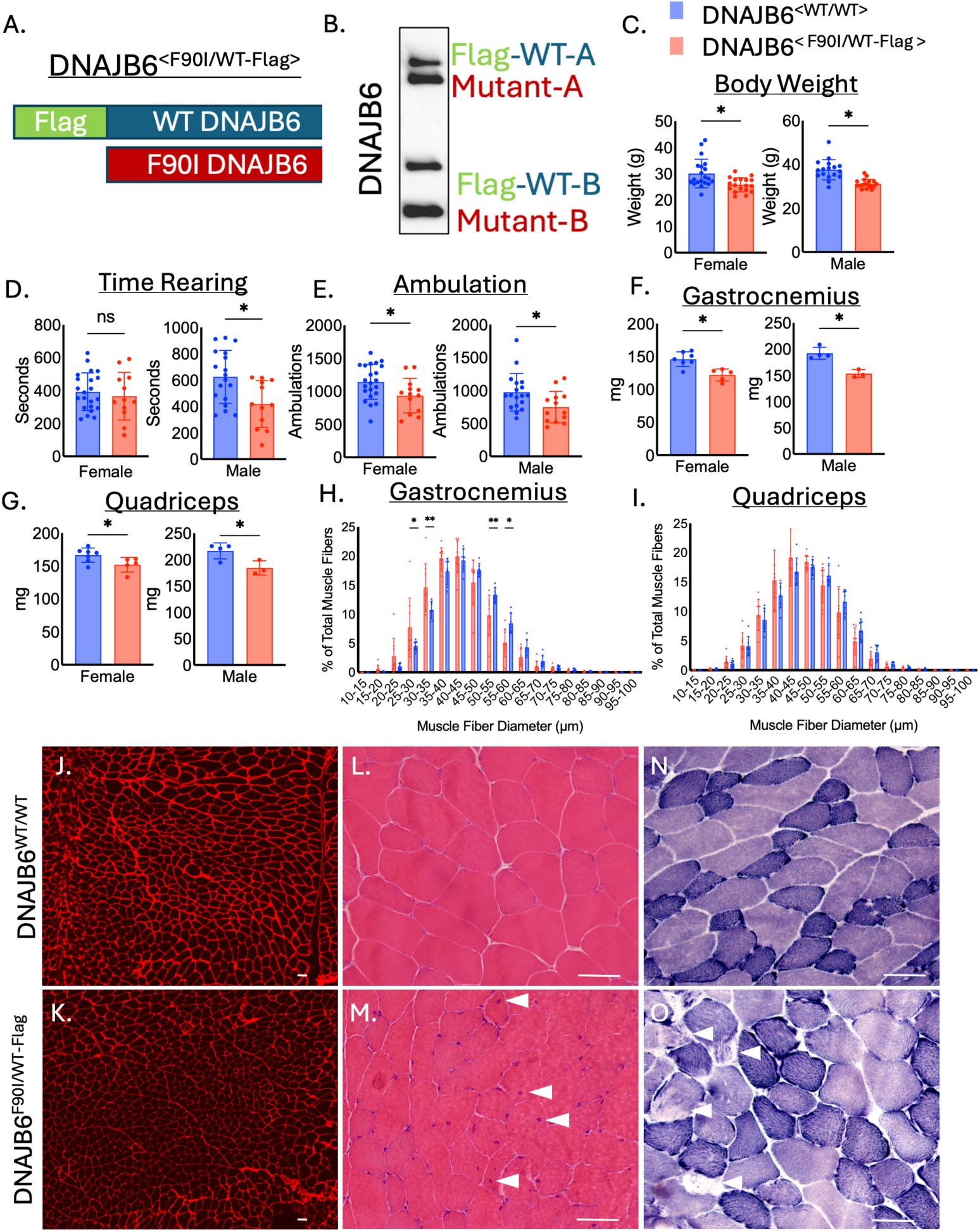
Flag-LGMDD1 mice permit allelic resolution of DNAJB6 and recapitulate features of LGMDD1. (A) Schematic of the *Dnajb6*^F90I/WT-Flag^ genotype, in which the wild-type allele contains an N-terminal 3×FLAG tag. (B) Representative anti-DNAJB6 immunoblot showing FLAG-tagged wild-type and untagged F90I DNAJB6a and DNAJB6b. (C–G) Body weight (C), time spent rearing during a 60-min activity-chamber assessment (D), total ambulations (E), and gastrocnemius (F) and quadriceps (G) muscle mass in 12-month-old female and male *Dnajb6*^WT/WT^ and *Dnajb6*^F90I/WT-Flag^ mice. (H and I) Gastrocnemius (H) and quadriceps (I) fiber-diameter distributions in 12-month-old female mice. (J and K) Representative laminin-stained gastrocnemius cross-sections from 12-month-old female *Dnajb6*^WT/WT^ and *Dnajb6*^F90I/WT-Flag^ mice. (L–O) Representative H&E (L and M) and NADH-tetrazolium reductase (N and O) staining of gastrocnemius muscle from 12-month-old female *Dnajb6*^WT/WT^ (L and N) and *Dnajb6*^F90I/WT-Flag^ (M and O) mice. Arrowheads indicate internal nuclei in (M) and fibers with altered internal architecture in (O). Scale bars, 50 μm. Data are presented as mean ± SD; each point represents one mouse. Comparisons in (C)–(G) were performed using multiple unpaired two-tailed *t* tests with Holm–Šídák correction. Fiber-diameter distributions in (H) and (I) were analyzed by two-way ANOVA with Šídák correction for multiple comparisons. \**P* < 0.05, \*\**P* < 0.01; ns, not significant. See also Figure S1.

At 12 months of age, female and male Flag-LGMDD1 mice had modestly reduced body weight compared with *Dnajb6* ^WT/WT^ controls (Fig. 1C). Male Flag-LGMDD1 mice spent less time rearing, whereas no significant difference was observed in female mice. Total ambulation was reduced in both sexes (Fig. 1D,E).

Gastrocnemius and quadriceps muscle masses were also reduced in female and male Flag-LGMDD1 mice (Fig. 1F,G). Gastrocnemius fiber-diameter distributions showed a modest shift toward smaller fibers (Fig. 1H, J, K). A similar pattern was observed in quadriceps, although the differences did not reach significance (Fig. 1I). Qualitative examination of H&E-stained gastrocnemius muscle showed mild myopathic changes, including fiber-size variability and internal nuclei (Fig. 1L,M). NADH-tetrazolium reductase (NADH-TR) staining identified scattered fibers with altered internal architecture, suggestive of myofibrillar disorganization (Fig. 1N,O). Overall, these findings recapitulate the phenotype previously reported in untagged *Dnajb6* ^F90I/WT^ mice.^8^

### Inhibition of protein degradation unmasks altered protein handling in LGMDD1 muscle

Immunoblot analysis of quadriceps muscle from 12-month-old *Dnajb6*^F90I/WT-Flag^ mice showed increased DNAJB6a, whereas DNAJB6b showed a nonsignificant increase compared with controls (Fig. 2A,B). Within Flag-LGMDD1 muscle, untagged F90I DNAJB6 also appeared more abundant than FLAG-WT DNAJB6. We previously found that *Dnajb6* transcript levels were unchanged in the untagged F90I knock-in model despite increased DNAJB6 protein, suggesting that DNAJB6 accumulates through a post-transcriptional mechanism.^1^ Flag-LGMDD1 muscle also contained increased polyubiquitinated proteins and desmin in whole-cell lysates, and increased HSP70 and p62 within the detergent-insoluble fraction (Fig. 2A,B). We additionally examined DNAJA4, a class A HSP40 co-chaperone identified in the proteomic analyses described below (Fig. 5), and found it increased in both fractions. Together, these findings indicate that Flag-LGMDD1 muscle exhibits basal alterations in protein homeostasis.

**Figure 2.**
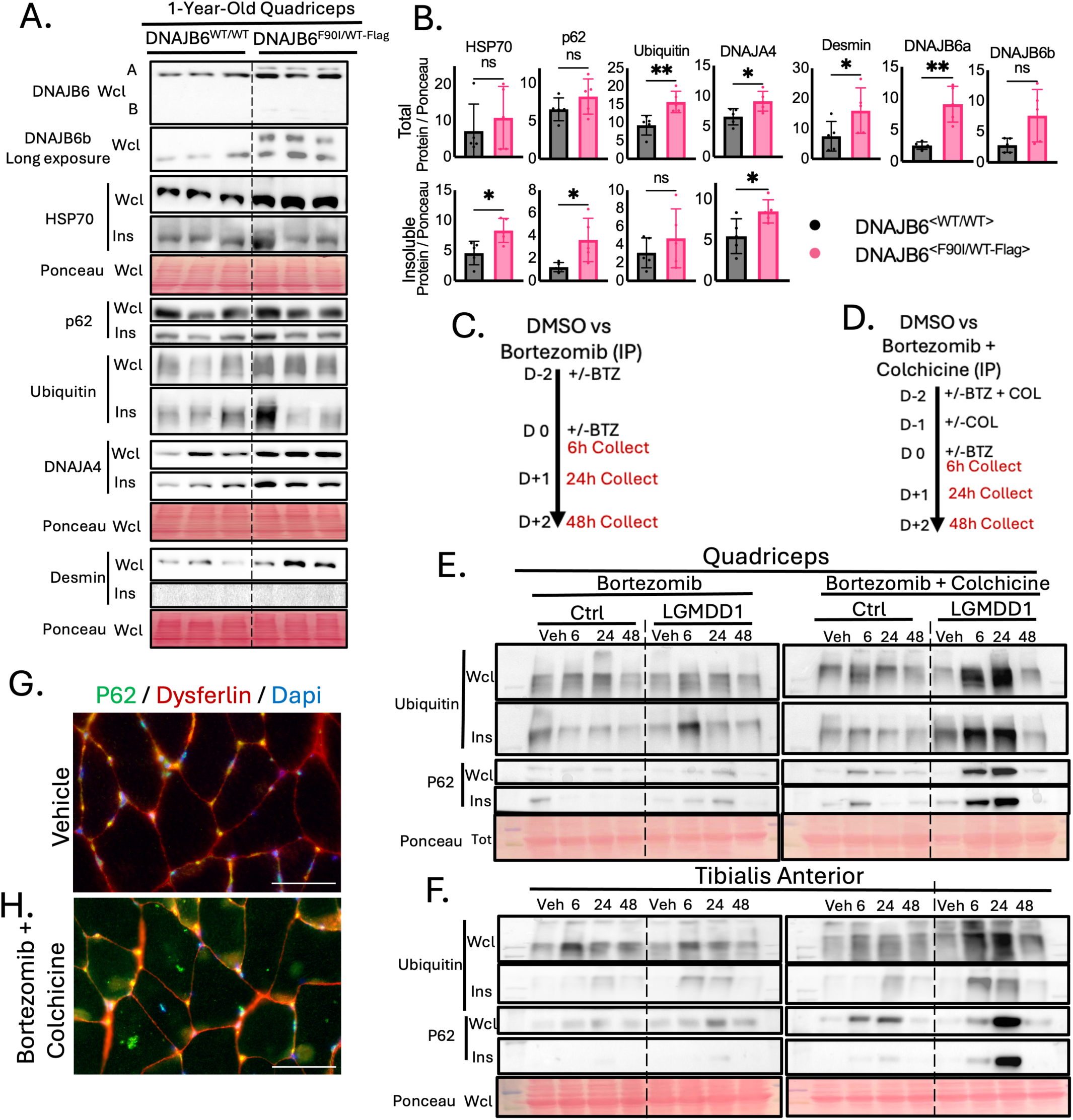
Biochemical characterization of proteostasis in LGMDD1 skeletal muscle (A) Immunoblot analysis of the indicated proteins in whole-cell lysates and detergent-insoluble fractions from quadriceps muscle of 12-month-old *Dnajb6^WT/WT^*and *Dnajb6^F90I/WT-Flag^* mice. Ponceau S staining was used to assess protein loading. (B) Quantification of the immunoblots in (A). Data are presented as mean ± SD; each point represents one mouse. Comparisons were performed using multiple unpaired two-tailed *t* tests with Holm–Šídák correction. (C and D) Treatment and tissue-collection schedules for bortezomib alone (C) or combined bortezomib and colchicine treatment (D). Six-to eight-month-old untagged *Dnajb6*^WT/WT^ and *Dnajb6*^F90I/WT^ mice received bortezomib (1 mg/kg), colchicine (0.4 mg/kg), or vehicle (DMSO) by intraperitoneal injection. For bortezomib treatment alone, mice received bortezomib on days −2 and 0. For combined treatment, mice received bortezomib and colchicine on day −2, colchicine on day −1, and bortezomib on day 0. Tissues were collected 6, 24, or 48 h after the final injection. (E and F) Immunoblot analysis of whole-cell lysates and detergent-insoluble fractions from quadriceps (E) and tibialis anterior (F) muscles of untagged *Dnajb6*^WT/WT^ and *Dnajb6*^F90I/WT^ mice treated according to the schedules in (C) and (D). Ponceau S staining was used to assess protein loading. A single time-course experiment is shown. Each lane represents muscle from a separate mouse collected at the indicated time point, and no inferential statistical comparisons were performed. (G and H) Representative immunofluorescence images of quadriceps from untagged *Dnajb6*^F90I/WT^ mice following vehicle (G) or combined bortezomib and colchicine treatment (H). Sections were stained for p62 (green), dysferlin (red), and DAPI (blue). Scale bars, 50 μm. BTZ, bortezomib; COL, colchicine; INS, detergent-insoluble fraction; Veh, vehicle; WCL, whole-cell lysate. \**P* < 0.05, \*\**P* < 0.01; ns, not significant.

Because DNAJB6 and HSP70 help maintain protein homeostasis, we next tested how LGMDD1 muscle responds when protein degradation is acutely interrupted. Six-to eight-month-old untagged *Dnajb6*^WT/F90I^ mice and *Dnajb6*^WT/WT^ controls were treated with bortezomib to inhibit the ubiquitin–proteasome system or with bortezomib and colchicine, a microtubule depolarizing agent that blocks autophagosome maturation to autolysosomes, to inhibit both proteasomal and autophagic degradation (Fig. 2C,D).^12^ Quadriceps and tibialis anterior muscles were collected 6, 24, or 48 h after treatment and analyzed by immunoblotting (Fig. 2E,F).

Bortezomib treatment increased polyubiquitinated proteins and p62 in both control and LGMDD1 muscle. Accumulation within the detergent-insoluble fraction appeared more prominent in LGMDD1 muscle and was observed in both quadriceps and tibialis anterior muscles (Fig. 2E,F). Combined bortezomib and colchicine treatment produced a more pronounced but transient accumulation of polyubiquitinated proteins and p62, particularly within detergent-insoluble fractions from LGMDD1 muscle. Combined treatment also produced prominent p62-positive puncta in LGMDD1 muscle compared with vehicle treatment, consistent with the accumulation of detergent-insoluble p62 observed by immunoblotting (Fig. 2G,H). These findings demonstrate acute interruption of proteasomal and autophagic degradation reveals altered protein handling in LGMDD1 muscle.

### Primary Flag-LGMDD1 myotubes retain altered DNAJB6 turnover and protein-handling abnormalities

To establish an in vitro system for evaluating allele-specific therapeutics, we generated primary myoblast cultures from *Dnajb6* ^WT/WT-Flag^ and *Dnajb6* ^F90I/WT-Flag^ mice. After 3 days of differentiation, both genotypes formed multinucleated myotubes with similar fusion indices, indicating that the F90I mutation did not produce an overt defect in myotube formation under these conditions (Fig. 3A,B). We next tested whether Flag-LGMDD1 myotubes reproduced the altered response to proteasome inhibition observed in skeletal muscle. Six-day differentiated myotubes were treated with MG132 for 4 h and collected immediately after treatment or at several time points following inhibitor washout (Fig. 3C). MG132 produced accumulation of polyubiquitinated proteins, p62, and HSP70 in whole-cell lysates and detergent-insoluble fractions. This accumulation appeared more prominent in Flag-LGMDD1 myotubes at several time points (Fig. 3D). Following MG132 removal, much of the accumulated material declined over the subsequent 48 h. At baseline, DNAJB6 also appeared more abundant in Flag-LGMDD1 myotubes than in controls, with F90I DNAJB6 more abundant than FLAG-WT DNAJB6 (Fig. 3D). This qualitative time course suggests that Flag-LGMDD1 myotubes reproduce the altered protein handling observed in skeletal muscle while retaining the ability to clear much of the accumulated material after proteasome inhibitor removal.

**Figure 3.**
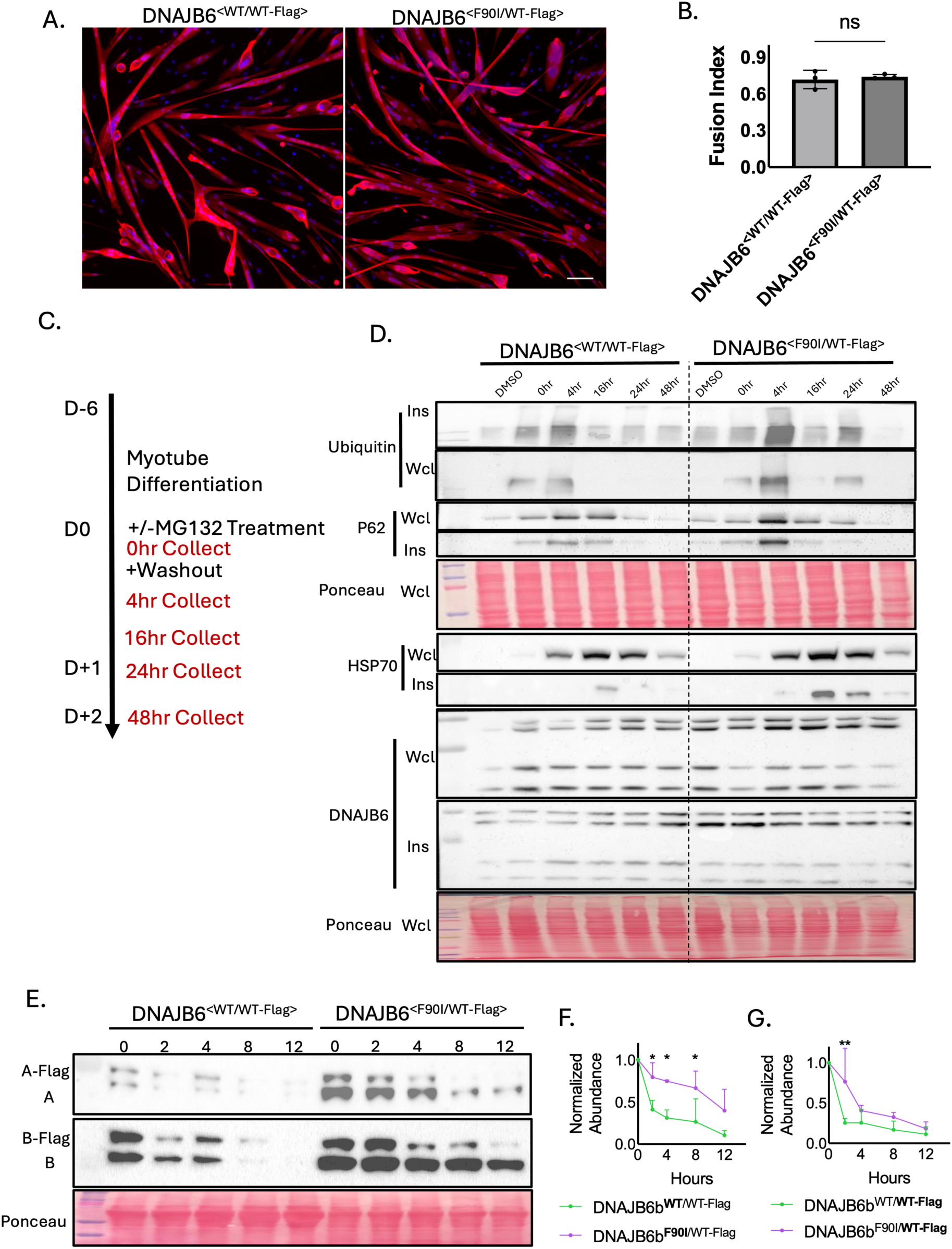
Generation and biochemical characterization of primary Flag-LGMDD1 myotubes (A) Representative immunofluorescence images of primary *Dnajb6*^WT/WT-Flag^ and *Dnajb6* ^F90I/WT-Flag^ myotubes after 3 days of differentiation. Myotubes were stained for α-actinin (red), and nuclei were labeled with DAPI (blue). Scale bars, 50 μm. (B) Fusion-index quantification. Data are presented as mean ± SD. Comparisons were performed using a two-tailed *t* test. ns, not significant. (C) MG132 treatment and washout schedule. Primary myotubes were differentiated for 6 days and treated with 10 μM MG132 or DMSO for 4 h. Cultures were collected immediately after treatment or following MG132 washout and recovery for 4, 16, 24, or 48 h. (D) Immunoblot analysis of whole-cell lysates and detergent-insoluble fractions from *Dnajb6* ^WT/WT-Flag^ and *Dnajb6* ^F90I/WT-Flag^ myotubes collected according to the schedule in (C). Ponceau S staining was used to assess protein loading. A single MG132 washout time course is shown. Each lane represents a separate culture collected at the indicated time point, and no inferential statistical comparisons were performed. (E) Representative anti-DNAJB6 immunoblot of *Dnajb6* ^WT/WT-Flag^ and *Dnajb6* ^F90I/WT-Flag^ myoblasts collected 0, 2, 4, 8, or 12 h after cycloheximide treatment. (F and G) Densitometric analysis of DNAJB6b following cycloheximide treatment. (F) Comparison of the untagged **WT** and **F90I** alleles. (G) Comparison of the **WT-Flag** allele in the *Dnajb6* ^WT/WT-Flag^ and *Dnajb6* ^F90I/WT-Flag^ backgrounds. Bold type in the graph keys indicates the allele quantified. DNAJB6b abundance was normalized to Ponceau S staining and expressed relative to the corresponding 0-h value. Data are presented as mean ± SD from three independent experiments and were analyzed by repeated-measures two-way ANOVA with Šídák correction for multiple comparisons using a single pooled variance. DMSO, vehicle control; INS, detergent-insoluble fraction; WCL, whole-cell lysate. \**P* < 0.05, \*\**P* < 0.01.

We next examined whether the increased abundance of F90I DNAJB6 resulted from altered protein stability. Myoblasts were treated with cycloheximide to inhibit new protein synthesis, and DNAJB6 abundance was measured over 12 h (Fig. 3E–G). Because of differences in isoform expression and stability, we focused on DNAJB6b. Untagged F90I DNAJB6b declined more slowly than untagged WT DNAJB6b (Fig. 3E,F). FLAG-WT DNAJB6b also declined slightly more slowly in *Dnajb6* ^F90I/WT-Flag^ myoblasts than in *Dnajb6* ^WT/WT-Flag^ myoblasts (Fig. 3E,G). These findings indicate that F90I DNAJB6b is degraded more slowly than WT DNAJB6b and suggest that the presence of the mutant allele may also alter the turnover of WT DNAJB6b in trans.^2,4^

### Systematic mismatch screening identifies an siRNA with greater than 250-fold selectivity for F90I DNAJB6

To identify siRNAs capable of selectively reducing F90I DNAJB6 while sparing the WT allele, we designed a series of 22-nucleotide siRNAs that tiled across the F90I variant (Fig. 4A). Each siRNA was fully complementary to the F90I allele and contained a single mismatch relative to the WT allele, with the position of the mismatch varying across the sequence. We selected 22-nucleotide siRNAs because the corresponding 19- and 21-nucleotide sequences had substantially more predicted off-target sites containing three or fewer mismatches (Table S1). Each siRNA was transfected into *Dnajb6* ^F90I/WT-Flag^ myoblasts at an initial screening concentration of 10 nM, and DNAJB6 protein abundance was measured three days later. The FLAG tag allowed simultaneous measurement of the WT and mutant DNAJB6 by immunoblotting without requiring separate allele-specific RT-PCR reactions (Fig. 4B). siRNAs 11 and 13 produced the greatest reduction in total DNAJB6, regardless of which allele, and were selected for dose-response testing (Fig. 4C). The two siRNAs had similar IC₅₀ values for total DNAJB6, whereas siRNA 11 had a lower IC₅₀ for F90I DNAJB6. siRNA 11 was therefore selected as the starting sequence for further optimization (Fig. 4D–F).

**Figure 4.**
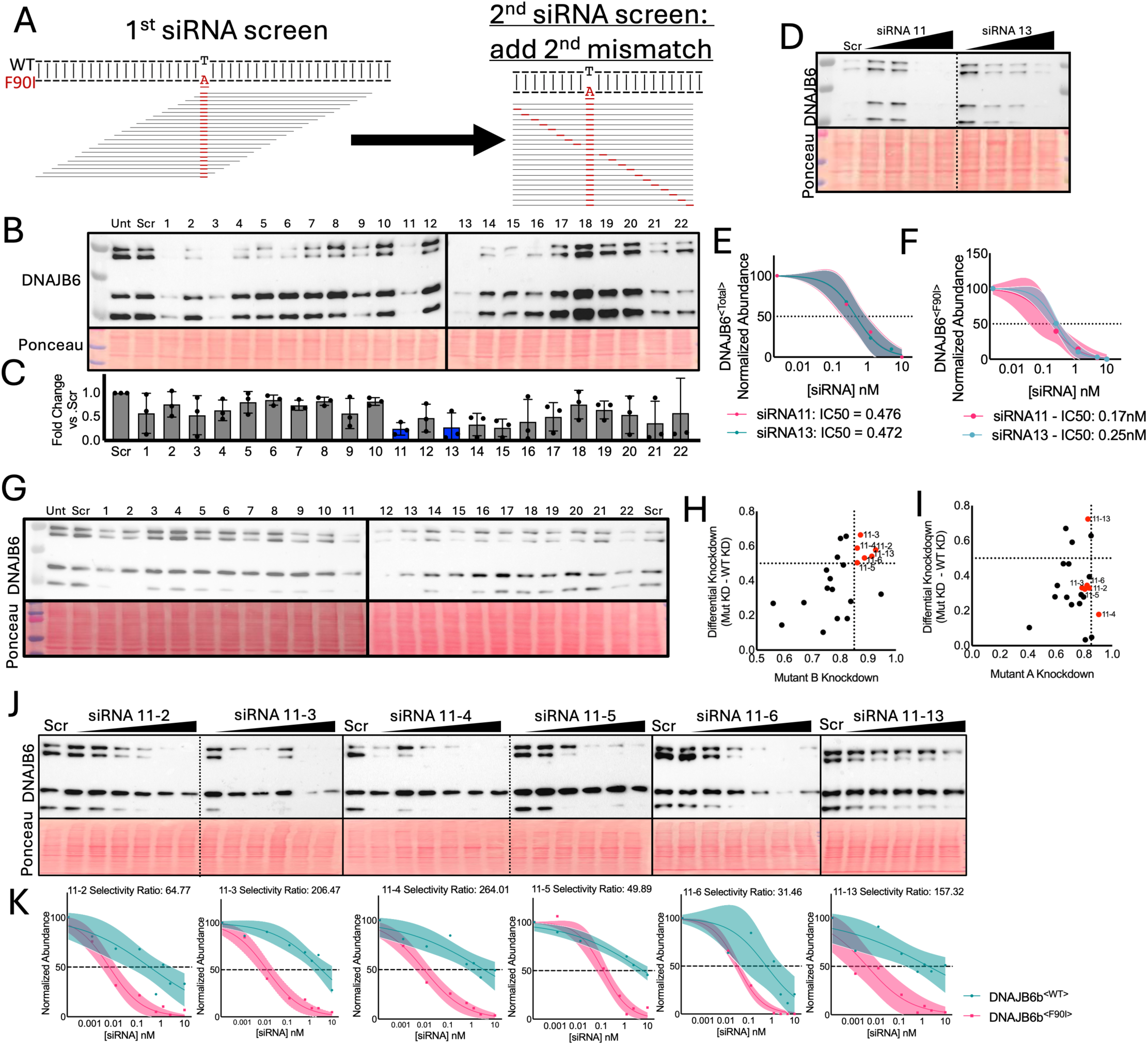
Identification and characterization of F90I-selective siRNAs (A) Two-stage siRNA screening strategy. First-generation 22-nucleotide siRNAs tiled across the F90I variant and were fully complementary to F90I Dnajb6, with a single mismatch relative to the WT allele. Second-generation siRNAs were derived from siRNA 11 by introducing an additional mismatch at successive positions across the sequence. (B and C) First-generation screen in *Dnajb6 ^F90I/WT-Flag^* myoblasts transfected with 10 nM siRNA and collected 3 days later. A representative anti-DNAJB6 immunoblot is shown in (B), and total DNAJB6 abundance relative to scrambled control siRNA is quantified in (C). siRNAs 11 and 13 are highlighted in blue. (D–F) Dose-response analysis of first-generation candidates siRNA 11 and siRNA 13 over a concentration range of 0.25–10 nM. A representative anti-DNAJB6 immunoblot is shown in (D). Dose-response curves for total DNAJB6 and F90I DNAJB6 are shown in (E) and (F), respectively. (G–I) Second-generation siRNA screen performed at 10 nM. A representative anti-DNAJB6 immunoblot is shown in (G). Mean F90I knockdown was plotted against differential knockdown for DNAJB6b (H) and DNAJB6a (I). Differential knockdown was calculated as the percentage knockdown of F90I DNAJB6 minus the percentage knockdown of FLAG-WT DNAJB6. The six candidates selected for further dose-response analysis are shown in red. (J and K) Dose-response analysis of six second-generation candidates over a concentration range of 0.000125–10 nM. Representative anti-DNAJB6 immunoblots are shown in (J), and DNAJB6b dose-response curves are shown in (K). DNAJB6 abundance was normalized to Ponceau S staining, with abundance following scrambled control treatment set to 100%. Observed values above 100% were retained without adjustment when calculating knockdown and differential knockdown. IC₅₀ values were calculated using unweighted least-squares nonlinear regression with a variable slope and the top and bottom constrained to 100% and 0%, respectively. Curves were fitted to the mean response at each concentration, and shaded regions indicate 95% confidence bands. The IC₅₀ selectivity ratio was calculated as the FLAG-WT IC₅₀ divided by the F90I IC₅₀. Data represent three independent experiments in (B)–(F), four independent experiments in (G)–(I), and three independent experiments in (J) and (K). Scr, scrambled control siRNA; Unt, untreated. See also Tables S1 and S2.

To improve allelic discrimination, we generated a second series of siRNAs by introducing one additional mismatch at successive positions across siRNA 11 (Fig. 4A). Each second-generation siRNA therefore contained one mismatch relative to the F90I allele and two mismatches relative to the WT allele. These siRNAs were screened at 10 nM in *Dnajb6*^F90I/WT-Flag^ myoblasts and analyzed three days after transfection (Fig. 4G). For each siRNA, we plotted mean F90I knockdown against differential knockdown, defined as the percentage knockdown of F90I DNAJB6 minus the percentage knockdown of FLAG-WT DNAJB6 (Fig. 4H,I). This allowed candidates to be evaluated simultaneously for mutant-allele potency and allelic discrimination. Based on four independent screening experiments, we selected six siRNAs that produced robust F90I knockdown with limited reduction of FLAG-WT DNAJB6. Candidate selection was based on DNAJB6b because of its greater relevance to skeletal-muscle disease and to avoid confounding by differences in isoform expression and stability. The six candidates were then tested across multiple independent dose-response experiments, with DNAJB6b quantified to determine the IC₅₀ for each allele (Fig. 4J,K). Selectivity was defined as the ratio of the FLAG-WT IC₅₀ to the F90I IC₅₀. Three siRNAs demonstrated greater than 100-fold selectivity for F90I DNAJB6b. The lead candidate, siRNA 11-4, had a selectivity ratio of 264, indicating that approximately 264 times the siRNA concentration was required to achieve 50% knockdown of FLAG-WT DNAJB6b compared with F90I DNAJB6b. Computational analysis identified two predicted binding sites for siRNA 11-4 containing three mismatches, in addition to the wild-type Dnajb6 transcript, and no predicted off-target sites containing at least 15 contiguous complementary nucleotides (Table S2).

### Acute generation of LGMDD1-associated ΔGF DNAJB6 produces proteomic changes concordant with chronic F90I expression

To identify proteins that respond to pathogenic DNAJB6 activity and could serve as exploratory markers of treatment response, we performed proteomic analysis of primary myotubes from *Dnajb6*^WT/F90I^ and *Dnajb6*^WT/WT^ mice. This identified 1,105 high-confidence proteins with altered abundance in F90I myotubes at nominal P < 0.05 (Fig. 5A, Table S3). Because F90I is expressed throughout life, these changes could reflect ongoing mutant DNAJB6 activity or chronic adaptation to it. We therefore sought a second comparison in which pathogenic DNAJB6 was generated acutely. A splice-switching oligonucleotide cannot introduce a point substitution (eg. F90I), but the G/F domain is largely encoded by Dnajb6 exon 5, so inducing exon 5 skipping in wild-type myotubes produced an in-frame protein lacking the G/F domain (Fig. 5A,B). This ΔGF protein corresponds to an LGMDD1-associated variant that causes severe, early-onset disease.^13^ F90I and ΔGF differ in the extent to which the G/F domain is disrupted, but both relieve the autoinhibition that normally prevents J domain engagement of HSP70.^6^ The acute ΔGF model contained 517 proteins with altered abundance at nominal P < 0.05, including 226 shared with the F90I dataset and 291 unique to the ΔGF dataset (Fig. 5A).

Across the 1,105 proteins altered in the F90I dataset, abundance changes correlated with those in the ΔGF dataset (r = 0.63, P < 0.0001; Fig. 5C), and the 226 shared proteins correlated more strongly (r = 0.82, P < 0.0001). The two models therefore share a subset of proteomic changes despite differing in variant, extent of G/F domain disruption, and duration of exposure. Proteins increased in both included DNAJB6 itself and the chaperones HSPA1A/HSP70, HSPA8/HSC70, and the HSP40-family co-chaperone DNAJA4 (Fig. 5C).

To characterize the broader pathways represented by the shared signature, ontology analyses were performed separately for proteins increased and decreased in both datasets, using all proteins identified by proteomic analysis as the background. Shared proteins with increased abundance were enriched for protein and RNA catabolic processes, regulation of inclusion-body assembly, sarcomeric structures including the Z-disc and costamere, and chaperone pathways involving HSF1 and HSP90 (Fig. 5D). Proteins with decreased abundance were enriched for pathways related to oxidative phosphorylation, ATP synthesis, and chromatin (Fig. 5E). We used concordance between the two models to prioritize candidates, reasoning that a change seen in only one could be due to chronic adaptation, loss of the entire G/F domain, or oligonucleotide treatment itself. DNAJA4 showed the largest increase of any chaperone or co-chaperone altered in both datasets and was selected for further evaluation.

### Allele-selective knockdown of F90I DNAJB6 reduces proteostatic abnormalities in Flag-LGMDD1 myotubes

To determine whether allele-selective knockdown improves downstream effects of mutant DNAJB6, Flag-control and Flag-LGMDD1 myoblasts were transfected with scrambled siRNA or siRNA 11-4 at 2.5 or 5 nM and differentiated for 3 days. Heat shock was used to increase protein unfolding, aggregation, and cellular demand on the HSP70 chaperone network, thereby revealing otherwise mild differences between control and Flag-LGMDD1 myotubes. Cultures were collected before heat shock or after 3 or 72 h of recovery from a 2-h heat shock at 42°C. Whole-cell lysates and detergent-insoluble pellet fractions were analyzed for markers of protein accumulation and aggregation, chaperone responses, and proteins associated with unfolded protein response signaling (Fig. 6A).

**Figure 5.**
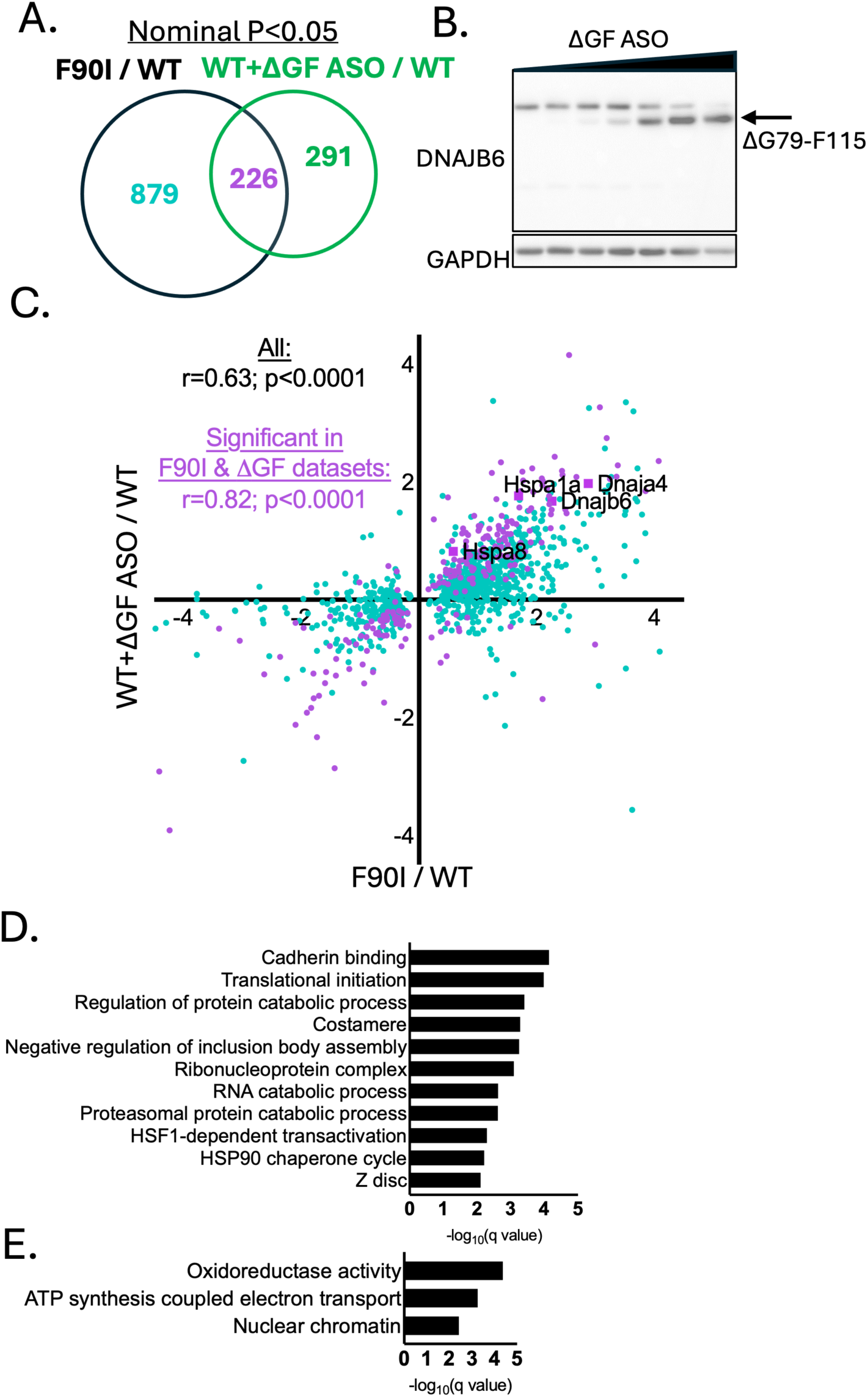
Proteomic comparison of F90I and acute ASO-induced ΔGF DNAJB6 myotubes. (A) Overlap between proteins with altered abundance in *Dnajb6 ^WT/F90I^* compared with *Dnajb6 ^WT/WT^* myotubes and in *Dnajb6 ^WT/WT^* myotubes treated with the ΔGF ASO compared with control-treated WT myotubes. ΔGF denotes the exon 5-skipping ASO that generates ΔG79–F115 DNAJB6. Only proteins identified at a 1% protein-level false discovery rate were included, with differential abundance defined at nominal *P* < 0.05. A total of 1,105 proteins were altered in F90I myotubes, 517 were altered following acute generation of ΔGF DNAJB6, and 226 were shared between the datasets. (B) Representative anti-DNAJB6 immunoblot following treatment of WT myotubes with increasing concentrations of the ΔGF ASO. The arrow indicates the ΔG79– F115 DNAJB6 protein generated by exon 5 skipping. GAPDH was used as a loading control. (C) Pearson correlations of log₂ fold changes between the F90I and ΔGF ASO datasets. The analysis included all 1,063 proteins altered in F90I myotubes. Proteins altered only in the F90I dataset are shown in turquoise, and the 216 proteins altered in both datasets are shown in purple. Selected chaperone proteins are indicated. Each proteomic comparison included three biological replicates per condition. (D and E) Pathway overrepresentation analysis of proteins with increased (D) or decreased (E) abundance in both datasets. All proteins identified by proteomic analysis were used as the background. Enrichment significance is shown as −log₁₀(*q* value), with *q* < 0.01 considered significant. See also Table S3.

**Figure 6.**
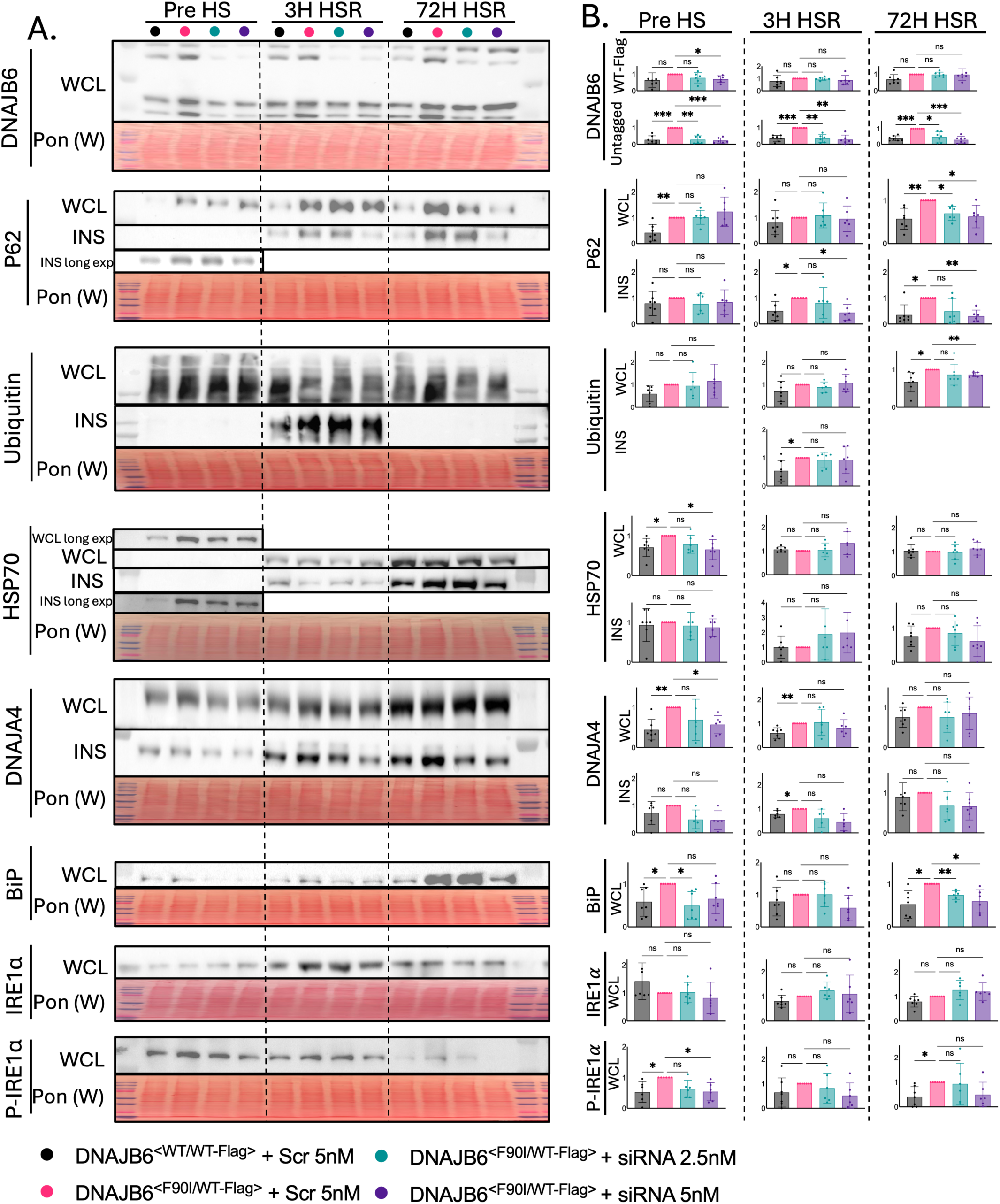
Effects of allele-selective F90I DNAJB6 knockdown on proteostatic stress in Flag-LGMDD1 myotubes. Near-confluent *Dnajb6^F90I/WT-Flag^* and *Dnajb6^WT/WT-Flag^* myoblasts were transfected with 5 nM scrambled siRNA or 2.5 or 5 nM siRNA 11-4 and differentiated for 3 days. Cultures were collected before heat shock or after a 2-hour heat shock at 42°C followed by 3 or 72 hours of recovery. (A) Immunoblot analysis of whole-cell lysates and detergent-insoluble fractions. Ponceau S staining was used to assess protein loading. (B) Densitometric quantification of the immunoblots in A. Within each time point, protein abundance was normalized to the scrambled siRNA-treated *Dnajb6^F90I/WT-Flag^*group, which was set to 1. This group was used as the common reference so that the genotype difference and the treatment effect could each be assessed against the same denominator within a time point. Data are presented as mean ± SD from four to six independent experiments. Statistical analyses were performed using repeated-measures one-way ANOVA with Geisser–Greenhouse correction, followed by Dunnett’s multiple comparisons test for three prespecified comparisons: *Dnajb6^WT/WT-Flag^* + Scr vs. *Dnajb6^F90I/WT-Flag^* + Scr, and *Dnajb6^F90I/WT-Flag^* + Scr vs. either 2.5 or 5 nM siRNA 11-4. No comparisons were made across time points. WCL, whole-cell lysate; INS, detergent-insoluble fraction; Pre HS, before heat shock; HSR, heat-shock recovery; Scr, scrambled control siRNA; ns, not significant. *P < 0.05, **P < 0.01, ***P < 0.001

Differences in markers associated with protein accumulation and aggregation were most apparent during recovery from heat shock. Flag-LGMDD1 myotubes showed increased p62 and ubiquitinated proteins following heat shock, particularly within detergent-insoluble fractions. The main baseline difference among these markers was increased whole-cell p62. Thus, heat-shock recovery provided a more sensitive setting for detecting abnormalities in protein aggregation.

In contrast, differences in chaperone abundance were most evident at baseline. Whole-cell HSP70 and DNAJA4 were increased in untreated Flag-LGMDD1 myotubes, consistent with the baseline proteomic findings. Heat shock produced a large increase in HSP70 abundance in both genotypes, possibly blunting the differences between control and Flag-LGMDD1 myotubes during recovery. DNAJA4 remained elevated in Flag-LGMDD1 myotubes after 3 h of recovery, but most other genotype differences were no longer apparent. Flag-LGMDD1 myotubes also showed increased BiP and phosphorylated IRE1α at baseline and later during heat-shock recovery, identifying additional differences in markers associated with unfolded protein response signaling.

Treatment with the lead siRNA selectively reduced F90I DNAJB6, with a small reduction in FLAG-WT DNAJB6 at some doses and time points (Fig. 6A,B). Allele-selective knockdown improved several of the abnormalities identified under baseline and heat-shock conditions. These included reduced accumulation of insoluble p62 during heat-shock recovery, reduced total ubiquitinated protein at 72 h of recovery, reduced baseline chaperone abundance, and reductions in the unfolded protein response (UPR)-associated proteins BiP and phosphorylated IRE1α. Although individual effects varied by dose and time point, the 5 nM dose produced the most consistent response. Overall, these experiments identify complementary molecular features of Flag-LGMDD1 myotubes: baseline differences in chaperone abundance and stress-dependent abnormalities in protein handling. Selective reduction of F90I DNAJB6 shifted features of both toward control values, providing proof of concept that allele-specific knockdown can lessen downstream effects of mutant DNAJB6.

## DISCUSSION

In this study, we developed an allele-resolved model of LGMDD1 and used it to test whether selective suppression of mutant DNAJB6 can improve downstream molecular features of disease. The FLAG-tagged wild-type allele allowed endogenous wild-type and mutant DNAJB6 to be measured within the same sample, while Flag-LGMDD1 mice recapitulated the functional, myopathological, and biochemical features of the previously characterized F90I knock-in model.^8^ Using this system, we developed siRNAs with strong discrimination between the two alleles. The lead candidate selectively reduced F90I DNAJB6 while largely preserving wild-type protein and improved several exploratory molecular features of LGMDD1 in primary myotubes. These findings provide cellular proof of concept for allele-specific knockdown as a potential treatment for LGMDD1.

Allelic resolution also allowed us to compare the abundance and turnover of wild-type and mutant DNAJB6. F90I DNAJB6 was more abundant than FLAG-WT DNAJB6 in skeletal muscle and declined more slowly following inhibition of protein synthesis in primary myoblasts. This is in agreement with our prior observation that DNAJB6 protein, but not transcript, is increased in the untagged F90I model and with recent evidence that LGMDD1-associated DNAJB6 mutants undergo reduced ubiquitination and degradation.^4,8,14^ FLAG-WT DNAJB6 also declined somewhat more slowly in cells containing F90I DNAJB6 than in control cells, consistent with earlier work reporting prolonged stability of both mutant and wild-type DNAJB6 in the presence of an LGMDD1 mutation.^4^ Because DNAJB6 forms multimeric complexes, mutant DNAJB6 could affect the turnover of wild-type DNAJB6 in trans, although an indirect effect on DNAJB6 turnover is also possible. This observation has one practical consequence for measuring knockdown selectivity. If the mutant allele slows wild-type turnover, then reducing F90I DNAJB6 may partly normalize the turnover of wild-type protein. A small decrease in FLAG-WT DNAJB6 following treatment, as we observed at baseline, could therefore reflect this indirect effect rather than loss of allelic discrimination.

Phenotypic characterization of endogenous LGMDD1 primary myotubes has been limited, so we first sought molecular phenotypes that could be measured reproducibly. In our prior study, F90I myotubes showed no overt abnormalities in proliferation, differentiation, or myofibrillar organization, and measurable differences were largely limited to a proteomic signature involving protein homeostasis and myofibrillar proteins, including increased HSP70.^1^ We therefore examined proteostasis in LGMDD1 skeletal muscle and primary myotubes at baseline and following two complementary proteostatic challenges. Following acute interruption of protein degradation, LGMDD1 muscle and myotubes accumulated more p62 and ubiquitinated proteins than controls, particularly within detergent-insoluble fractions. Both total and insoluble material declined after inhibitor removal, suggesting that protein degradation is not broadly impaired by LGMDD1 mutations. These findings could reflect a greater substrate load entering degradative pathways, but flux was not directly measured and the basis of insoluble protein accumulation remains unclear. These experiments were single time courses analyzed qualitatively and were not subjected to statistical comparison.

Prior work indicates that LGMDD1 mutations allow DNAJB6 to engage and stimulate HSP70 inappropriately, reducing HSP70 availability for other quality control functions.^6–8^ We did not measure HSP70 availability, but our findings can be interpreted within this model. At baseline, Flag-LGMDD1 myotubes showed increased HSP70, DNAJA4, whole-cell p62, BiP, and phosphorylated IRE1α, without accumulation of detergent-insoluble material. One explanation is that mutant DNAJB6 imposes a low level of ongoing proteostatic stress that cells offset by increasing chaperone abundance, and that this compensation is sufficient to prevent aggregation under basal conditions. Following heat shock, the differences between genotypes changed character rather than simply increasing. Chaperone differences were largely lost at 3 h of recovery, when heat shock increased HSP70 in both genotypes, while p62 and ubiquitinated proteins accumulated within the detergent-insoluble fraction, a change not seen at baseline. DNAJA4 and HSP70 were also recovered in the insoluble fraction during recovery. This pattern suggests that the capacity to compensate is limited and can be exceeded when additional unfolded protein load is imposed. Other explanations for the increased baseline chaperone abundance are possible. Because mutant DNAJB6 is degraded more slowly and forms multimeric complexes, increased DNAJA4 and HSP70 may be due to co-stabilization of complex members rather than increased synthesis, and both proteins were also recovered in the detergent-insoluble fraction, where they may be less accessible to degradation. Distinguishing transcriptional induction from altered protein turnover would require measurement of the corresponding transcripts, which we did not perform. Selective reduction of F90I DNAJB6 lowered baseline chaperone and UPR marker abundance, reduced insoluble p62 accumulation during heat-shock recovery, and reduced total ubiquitinated protein at 72 h of recovery, indicating that these features require continued expression of the mutant allele.

The proteomic comparison between chronic F90I expression and acute generation of ΔGF DNAJB6 provided another approach to identify proteins responsive to pathogenic DNAJB6 activity. Despite differences in the specific variant and exposure duration, the two models produced concordant proteomic changes. These alterations included increased DNAJB6, HSP70, HSC70, and DNAJA4, alongside shifts in other proteins related to protein homeostasis, sarcomeric structure, and energy metabolism. Enriched terms included HSF1-dependent transactivation and the HSP90 chaperone cycle, the latter reflecting increased abundance of the HSP90 co-chaperones AHSA1 and SUGT1.^15^ DNAJA4 showed the largest increase of any chaperone or co-chaperone altered in both comparisons, exceeding the increase in DNAJB6 itself, and we confirmed increased DNAJA4 in Flag-LGMDD1 myotubes and skeletal muscle by immunoblot. DNAJA4 also decreased following allele-selective knockdown. DNAJA4 is a transcriptional target of HSF1 and is induced by proteotoxic stress, so its increase may reflect activation of the heat shock response rather than a specific relationship with mutant DNAJB6.^16,17^ Whether DNAJA4 contributes to LGMDD1 pathology is unknown. Few proteins in either comparison reached significance after correction for multiple comparisons, and DNAJA4 did not. We therefore applied a nominal threshold, and the broader shared signature and ontology findings should be considered exploratory. Reproducibility across two independent proteomic comparisons and confirmation by immunoblot in muscle and myotubes are the main reasons we prioritized DNAJA4 for further evaluation.

Flag-LGMDD1 myotubes also showed increased BiP and phosphorylated IRE1α at baseline and during heat-shock recovery, and both decreased following allele-selective knockdown. Altered DNAJB6 function has been linked to activation of ER stress markers in zebrafish, and DNAJB6 has been shown to partially localize to the ER following tunicamycin treatment.^18^ We measured protein abundance only and did not evaluate downstream signaling, so these results do not establish that the unfolded protein response is activated. Cytosolic proteostatic stress can influence ER stress signaling indirectly, and in skeletal muscle the sarcoplasmic reticulum is continuous with the ER, which may provide additional routes by which altered cytosolic protein handling affects these markers. It is also possible that BiP and phosphorylated IRE1α increase as part of a broader stress response. BiP abundance was unchanged in both proteomic datasets and most other ER chaperones were not increased, so this observation requires further evaluation.

Preserving wild-type DNAJB6 expression while reducing the mutant allele is thought to be a critical safety requirement for LGMDD1 therapeutics. While complete loss of DNAJB6 is lethal in mice, haploinsufficiency is largely tolerated in mice, which do not develop skeletal muscle disease, suggesting that a successful treatment must maintain a threshold level of wild-type protein. For our siRNA optimization strategy, rather than selecting the most allele-selective first-generation sequence and then adding a second mismatch, which yielded poor potency and reproducibility in pilot testing, we prioritized maximum initial potency against the mutant allele as the baseline for systematic mismatch placement. This approach is similar to others’ allele-selective oligonucleotide development strategies.^19,20^ The resulting lead candidate demonstrated 264-fold selectivity for F90I DNAJB6b alongside a favorable predicted off-target profile. Screening directly at the protein level allowed both alleles to be quantified simultaneously. This direct measurement is critical because the altered stability of mutant DNAJB6 means that changes in transcript abundance may not fully predict the magnitude or timing of protein reduction.

The selectivity we observed in vitro does not by itself establish a therapeutic window. In allele-selective oligonucleotide programs for Huntington disease, the limiting factor proved to be potency in target tissue rather than allelic discrimination. First-generation compounds designed against SNPs in phase with the pathogenic allele showed allelic selectivity in vitro but were discontinued after producing insufficient lowering of mutant protein in patients, whereas a later compound incorporating modified backbone chemistry achieved substantial mutant protein reduction while preserving wild-type protein.^21–23^ Whether allele-selective knockdown can reduce mutant DNAJB6 sufficiently in skeletal muscle while preserving wild-type protein will require dose-ranging studies in vivo.

The siRNA described here targets the murine F90I sequence and is a tool for preclinical evaluation rather than a candidate human therapeutic. A human equivalent directed against F89I would apply only to individuals carrying that variant. For human translation, targeting a SNP in phase with the pathogenic allele would extend the approach to a larger proportion of patients, although not to all. This differs from isoform-selective knockdown of DNAJB6b, which we described previously and which is variant independent but reduces wild-type DNAJB6b along with mutant protein.^1^ The two strategies therefore trade breadth of applicability against preservation of wild-type protein.

Although these findings provide a proof of concept, several limitations warrant consideration. First, because of tissue availability, the in vivo characterization compared 12-month-old Flag-LGMDD1 mice with untagged wild-type controls rather than mice carrying a FLAG-tagged wild-type allele, and consequently a phenotypic contribution from the FLAG tag cannot be entirely excluded. This possibility is mitigated by the fact that the phenotype closely resembled that of the previously characterized untagged F90I model, and FLAG-tagged controls were used for the cell-culture experiments.^8^ Second, the proteomic and treatment experiments used different culture conditions. Myotubes for proteomic analysis were differentiated for four days before morpholino treatment and collected at day 10, whereas myoblasts for siRNA experiments were transfected at confluency and collected after three days of differentiation. Each proteomic comparison also included only three biological replicates. Differences between the two datasets are therefore expected, and we place greater confidence in changes observed by both approaches. Third, the conditions under which we delivered siRNA differ from those of a therapeutic given to patients in two respects, the state of the target cell and the method of delivery. The siRNA was introduced into near-confluent myoblasts before differentiation rather than into mature myotubes. All endpoints were evaluated after myotube formation, but confounding from differences in differentiation kinetics cannot be ruled out, and this approach does not reproduce treatment of established myofibers. It may instead partly model exposure during active regeneration, as occurs in diseased muscle. We also used unmodified siRNA delivered with a transfection reagent, which does not address delivery to skeletal muscle in vivo. Muscle-targeting oligonucleotide conjugate platforms developed for other neuromuscular diseases now achieve substantial delivery to skeletal muscle, and incorporating such a platform would be required to extend this work in vivo. Fourth, the treatment studies relied on a single lead siRNA, and its off-target profile was assessed computationally rather than experimentally. Transcriptome-wide or proteome-wide analysis following treatment would be needed to exclude sequence-dependent off-target effects. Finally, the observed improvements were limited to molecular markers rather than functional endpoints, which may not manifest in in vitro models of LGMDD1. Demonstrating correction of these disease-associated markers does not equate to restoration of muscle function. Adaptation to human LGMDD1 cell lines alongside in vivo evaluation of safety and functional benefit will be required to determine the therapeutic potential of this approach.

Allele-selective reduction of F90I DNAJB6 improved multiple baseline and stress-dependent molecular features of altered protein homeostasis in primary Flag-LGMDD1 myotubes. Whether preserving wild-type DNAJB6 confers a specific benefit over nonselective suppression remains to be established, as these experiments did not include a nonselective comparator. These results establish proof of concept for allele-specific knockdown and support continued translation of this strategy for LGMDD1.

## METHODS

### Animal and Experimental Protocols

All animal experimental protocols were approved by the Animal Studies Committee of Washington University School of Medicine. For the CRISPR/Cas9-mediated *knock in* of an N-terminal 3x Flag-tag immediately downstream of the 1^st^ ATG, we used two gRNAs (TCATTGGGCAGTGTTATCATNGG, GGCTGTATTTTTTTATTTGCNGG) and a donor oligonucleotide containing the 3x Flag-tag flanked by 60bp of homologous sequence to the targeted region. Zygotes were microinjected with a mixture of Cas9 mRNA, gRNAs, and oligos and transferred to pseudopregnant female mice. Genotyping of F0 born pups using tail-extracted DNA and primers flanking the Flag insertion site led to the identification of a founder mouse carrying the Flag-tag on one allele. Mice in the subsequent F1 generation who carried the Flag-tag were healthy, normal in size, and did not display any phenotype compared to their wild-type littermates. The generation of these mice was carried out by the genome engineering and stem cell core and the mouse genetics core facility at Washington University. Animal lines were bred on a C57BL/6 background (The Jackson Laboratory) to at least the F5 generation for phenotypic analysis. Control animals were *Dnajb6^WT^*^/WT-Flag^ or *Dnajb6*^WT/WT^ littermates as indicated. Mice were housed in a temperature-controlled environment with 12-hour light/12-hour dark cycles, in which they received food and water ad libitum. Mice were euthanized, and skeletal muscles were dissected. Muscle was flash frozen in liquid nitrogen-cooled isopentane and stored at −80°C. Mice received bortezomib (1 mg/kg), colchicine (0.4 mg/kg), or vehicle by intraperitoneal injection.

### Genotyping of mice

Genomic DNA was extracted from mouse tails using KAPA Express DNA extraction Kit (Kapabiosystems, KK7103) and standard PCR was performed using Lambda Biotech (206,811) and gene-specific primers. PCR products were subjected to electrophoresis on a 1.5% agarose gel. After traditional PCR verification of germline transmission of *Dnajb6*^WT-Flag^ alleles in F1 litters, mouse genotypes were determined by real-time PCR analysis (Transnetyx, Cordova, TN, USA) using tail biopsy samples.

### Animal behavioral tests

After habituation and handling in the Washington University Animal Behavior Core, mice were assessed for locomotor activity. All tests were conducted during the light phase of the light/dark cycle. Motor and exploratory behavior of mice were evaluated over a 1-hour period in transparent (47.6 cm by 25.4 cm by 20.6 cm high) polystyrene enclosures. Each cage was surrounded by a frame containing a 4 × 8 matrix of photocell pairs, the output of which was fed to an on-line computer (Hamilton-Kinder, LLC, Poway, CA). Variables that were analyzed included the total number of ambulations and time spent rearing on hindlimbs.

### Primary mouse myoblast isolation, culture, and treatment

Primary mouse myoblasts were isolated and cultured as previously described.^24^ Hindlimb muscles were isolated from *Dnajb6*^WT/WT-Flag^ and *Dnajb6*^F90I/WT-Flag^ mice, minced into small pieces, digested with 1 mL per muscle of enzyme mix (500 U/mL type II collagenase [MP biomed], 1.5 U/mL Collagenase D [Sigma] 2.5 U/mL Dispase II [Sigma], 2.5 mM CaCl2, PBS up to 1 mL), shook at 37°C for 1 hour, pelleted at 300xG for 5 minutes, and the supernatant was discarded. The pellet was resuspend in myoblast isolation medium (high-glucose Dulbecco’s Modified Eagle Medium [DMEM, Gibco], supplemented with 20% fetal bovine serum, [FBS, Gibco], 10% horse serum [HS, Gibco], 0.5% chicken embryo extract [CEE, MP Biomedicals], 2.5 ng/mL basic fibroblast growth factor [bFGF, PeproTech], 1% penicillin-streptomycin [Gibco], 10 µg/mL gentamicin [Gibco], 100 µg/mL kanamycin [Gibco], 1 µg/mL amphotericin B [Gibco], and 5 µg/mL Plasmocin prophylactic [Invivogen, San Diego, CA]), seeded on Matrigel-precoated dishes (0.7mg/mL) at 10-20% surface coverage, and incubated at 37°C and 5% CO2. After cellular outgrowth from muscle explants was observed, any remaining tissue chunks were transferred to new Matrigel coated plates for additional rounds of myoblast outgrowth. Prior to confluence, outgrown cells were detached using TrypLE Express and to improve myoblast purity, cells were pre-plated on collagen coated plates (type I rat tail collagen [CORNING] in sterile water at 0.1mg/mL) in myoblast isolation medium, for 1 hour at 37°C and 5% CO2. The supernatant was then transferred to Matrigel-coated dishes while cells on the collagen coated plates were discarded. Purified myoblasts were subsequently grown on 0.43mg/mL Matrigel-precoated dishes (Passages 1-3) or 0.2% gelatin-coated dishes (Passages 4-20) in proliferation medium (1:1 ratio of HG DMEM and Ham’s F10 nutrient mix, supplemented with 20% FBS, 10% horse serum, 1% penicillin-streptomycin, 100 µg/mL kanamycin, 10 µg/mL gentamicin, 5 µg/mL Plasmocin prophylactic, 1 µg/mL amphotericin B, and 2.5ng/mL FGF2), For differentiation into myotubes, myoblasts were seeded at a high baseline confluency in proliferation media, and media was infrequently changed to allow serum levels to be gradually consumed. Unmodified siRNAs with dTdT overhangs [Sigma] were transfected using endoporter [GeneTools] according to the manufacturers protocol.

For protein synthesis inhibition, myoblasts were treated with cycloheximide at 100µg/ml. Octaguanidine dendrimer-conjugated morpholino oligonucleotides (Gene Tools) were used in this study. ΔGF (5’ACGGACCAGGCAGGTACTTACCA -3’; Gene Tools) and negative control (5’-CCTCTTACCTCAGTTACAATTTATA-3’; Gene Tools). Primary myotube cultures were treated with morpholinos (0.375µM) after four days in culture, replenished at day 7, and collected at day 10 for mass spectrometry.

### siRNA design and off-target prediction

Candidate siRNA sequences were screened for potential binding sites using GGGenome (https://gggenome.dbcls.jp/), identifying sequences containing three or fewer mismatches relative to each siRNA. First-generation siRNAs were queried against both the mouse genome (GRCm39/mm39) and the RefSeq mouse RNA database; second-generation siRNAs were queried against the RefSeq mouse RNA database. Second-generation siRNAs were additionally queried using NCBI BLAST to identify transcripts containing at least 15 contiguous complementary nucleotides, a length at which off-target silencing has been reported.^25,26^ Predicted binding sites for first- and second-generation siRNAs are provided in Tables S1 and S2.

### Histochemistry

For histological analysis of skeletal muscle, samples were mounted in tragacanth gum (10% solution, Sigma- Aldrich, G1128), flash-frozen in a bath of isopentane suspended in liquid nitrogen and stored at − 80 °C until ready to be sectioned to an 8 μm thickness. For the H&E stain, a 1% aqueous solution of eosin Y (Sigma E-6003) was prepared in deionized water, and Harris hematoxylin stain (Lerner Laboratories, 1,931,382) was filtered before use. Slides in a metal staining rack were immersed in the filtered Harris hematoxylin for 10 s, then transferred to a beaker of tap water, and rinsed until the water was clear. Then, the slides were immersed in eosin stain for 30 s and again rinsed with tap water. Then, sections were dehydrated in ascending alcohol solutions (50%, 70%, 80%, 95% × 2, 100% × 2) and cleared with xylene three or four times, and a glass coverslip was mounted to the glass slide using Permount. For nicotinamide adenine dinucleotide tetrazolium reductase (NADH-TR) histochemistry, sections were incubated in a working solution of 0.05 M Tris buffer (pH 7.6) containing nitro blue tetrazolium (NBT) and NADH for 30 minutes at 37 °C. Following incubation, sections were rinsed with deionized water and differentiated through graded acetone solutions (30%, 60%, 90%, 60%, 30%) to remove unbound background dye.

### Immunofluorescence

Muscle sections were fixed using 3.7% PFA for 10 minutes, rinsed 3 times in 1x PBS, followed by additional fixation and partial permeabilization for 5 min in ice-cold acetone. The muscle sections were rinsed again 3 times in 1x PBS, then further permeabilized for 5 minutes in 0.5% Triton X-100 in 1x PBS at room temperature. Slides were rinsed again 3 times in 1x PBS, then blocked for 90 min at room temperature in 0.22µm filtered PerkinElmer blocking reagent (FP1012). Primary antibodies were diluted in blocking reagent and incubated at 4 °C overnight. After six rinses for 5 minutes each with 1 × PBS, appropriate fluorophore-conjugated secondary antibodies at 1:500 dilution and 5µg/mL 4′,6-diamidino-2-phenylindole (DAPI) were added to the slides in blocking reagent and incubated for 90 minutes at room temperature with light protection. Slides were rinsed with 1 × PBS again six times for 5 minutes each and mounted with a coverslip using Mowiol 4–88 (Sigma-Aldrich, 81,381) containing 1,4-diazabicyclo [2, 2, 2] octane (DABCO) antifading reagent. The following primary antibodies and dilutions were used for immunostaining: rabbit anti-laminin (abcam; ab11575; 1:500), mouse anti-dysferlin (Vector Laboratories; VP-D503; 1:250), rat anti-LAMP2 (abcam; ab13524; 1:200), rabbit anti-LC3B (Millipore Sigma; L7543; 1:200), rabbit anti-SQSTM1 (P62) (Proteintech; 18-420-1-AP; 1:1000), and mouse anti-Ubiquitin (P4D1) (Cell Signaling; 3936; 1:200). Secondary antibodies were the following: goat-anti-mouse Alexa555 (Invitrogen; A-21422), goat-anti-rabbit Alexa488 (Invitrogen; A-11008), goat-anti-rat Alexa647 (abcam; 150159).

Primary mouse myoblasts and myotubes were grown, stained, and imaged directly on cell culture plastic coated with Matrigel. All cells were washed 3 times with 1x PBS, fixed in 4% PFA for 10 minutes, washed 3 times again with 1x PBS, permeabilized with 0.5% Triton X-100 in PBS for 10 minutes, washed 3 times again in 1x PBS, and then blocked with 3% BSA in PBS for 2 hours at room temperature. Cells were stained with primary antibody rabbit anti-alpha-actinin (Abcam; ab52866; 1:300) in 3% BSA and 0.1% Saponin in PBS at 4°C overnight, followed by washing 3 times with PBS, and a 2 hour light protected room temperature incubation in goat-anti-rabbit Alexa555 secondary antibody (Invitrogen; A-21428; 1:1500) in 3% BSA in PBS containing 5µg/mL 4′,6-diamidino-2-phenylindole (DAPI). Cells were rinsed a final 3 times with PBS prior to imaging.

### Imaging and image analysis

H&E, NADH, P62, and Ubiquitin stained skeletal muscle sections were imaged using a Leica DM4 B microscope (Leica Microsystems, Germany), and images were acquired using Leica Application Suite X (LAS X) software. For quantitation of muscle fiber diameter, entire sections (3) of laminin-stained skeletal muscle were scanned using a Zeiss Axio Scan.Z1 microscope with a Plan-Apochromat 20X objective equipped with a Hamamatsu Orca Flash sCMOS camera and Zen 3.1 (blue edition) software. Quantification of the minimal feret diameter of myofibers was performed using the MuscleJ2.0 plugin installed in Fiji (ImageJ, National Institutes of Health, Bethesda, MD, USA). Muscle sections were reviewed qualitatively by a clinical neuromuscular pathologist for standard myopathic features, including fiber-size variability, internal nuclei, and abnormalities in internal architecture and myofibrillar organization. Differences between myotubes were quantitated as a ratio of nuclei within multinucleated alpha-actinin positive myotubes divided by the total number of nuclei. For each experimental replicate, nuclei were counted from 5 random fields taken with a 10X objective equipped in a NIKON Eclipse 80i fluorescence microscope.

### Western blot

Muscle tissues and cultured cells were homogenized using RIPA lysis buffer (50 mM Tris–HCl, pH 7.4, 150 mM NaCl, 1% NP-40, 0.25% Na-deoxycholate, and 1 mM EDTA) supplemented with protease inhibitor cocktail (Millipore Sigma), and lysates were centrifuged at 16,000g for 10 minutes. Protein concentrations were determined using a BCA Protein Assay Kit (Thermo Fisher Scientific) and measured with the BioTek Epoch plate reader at 562 nm. Aliquots of unfractionated, total cell lysates were solubilized in Laemmli sample buffer, and equal amounts of proteins were separated on 12% SDS-PAGE gels. Proteins were transferred onto nitrocellulose membranes and total protein transfer was assessed by Ponceau S staining. The membranes were subsequently washed with PBS containing 0.1% Tween-20 until the stain was completely removed, followed by blocking with 5% nonfat dry milk in PBS with 0.1% Tween-20 for 1 hour. The membrane was then incubated with primary antibodies, specific to the protein of interest, in 5% nonfat dry milk in PBS with 0.1% Tween overnight at 4°C. After incubation with the appropriate secondary antibody conjugated with HRP, Clarity™ Western ECL (Biorad) was used for protein detection. Immunoblots were obtained using the G:Box Chemi XT4, Genesys, version 1.1.2.0 (Syngene). The antibodies used were as follows: anti–rabbit GAPDH (Cell Signaling Technology; 2118), anti–rabbit desmin (Abcam; ab8592), anti–rabbit DNAJB6 (Abcam; ab198995), anti–mouse HSP70 (Enzo Life Sciences; ADI-SPA-812), anti–rabbit P62 “SQSTM1” (Proteintech; 18420-1-AP), anti–rabbit DNAJA4 (Invitrogen; MA5-52922), anti–mouse Ubiquitin (Enzo; ABS840), anti–rabbit IRE1 (Abcam; ab37073), anti–rabbit phospho-IRE1 alpha (Invitrogen; PA116927) and anti–mouse Bip/GRP78 (BD transduction labs; 610979). The following secondary antibodies were used: anti–mouse HRP (Cell Signaling Technology; 7076S), anti–rabbit HRP (Cell Signaling Technology; 7074S).

### Insoluble pellet assay

Tissue homogenates were precleared by centrifugation at 16000g for 15 min at 4°C, and an aliquot of the supernatant was collected and labeled the total fraction. The remaining supernatant was centrifuged at 100,000 g for 30 minutes at 4°C and the supernatant was discarded. The pellet was washed with 150µl of lysis buffer, sonicated on ice, and then centrifuged at 100,000 g for 30 minutes at 4°C. The washing step was repeated, and the resultant pellet (insoluble fraction) was resuspended in 7M urea buffer with 2M thiourea, 4% CHAPS and 30mM tris (pH 8.5) and used for subsequent analysis.

Cell lysates were precleared with a 30-second low-speed spin, and an aliquot of the supernatant was collected and labeled the total fraction. The remaining supernatant was centrifuged at 21,000g for 10 minutes at 4°C. The supernatant was discarded and the pellet was then washed with lysis buffer, sonicated on ice, then centrifuged at 21000 x g for 5 min at 4°C. The washing step was repeated and the resultant pellet (insoluble fraction) was resuspended in 7M Urea and 100U/mL Benzonase and incubated at 22-25°C for 15 minutes.

Samples were then centrifuged at 21000 x g at 4°C for 10 min. The supernatant is then collected and used for subsequent analysis.

### Mass spectrometry

Primary myotubes were grown, treated with morpholinos, and cell lysates were prepared as described above. The lysates were digested in an S-Trap micro spin column (Protifi, USA) according to the manufacturer’s instructions. After elution and dried, peptides were suspended in the LCMS-grade water containing 0.2% formic acid and 2% acetonitrile for further LC-MS/MS analysis. LC-MS/MS data were acquired in data-dependent acquisition (DDA) mode using an EASY-nLC 1200 (ThermoFisher Scientific, San Jose, CA) coupled to a Q Exactive HF hybrid quadrupole-Orbitrap mass spectrometer (ThermoFisher Scientific, San Jose, CA). Peptides were separated on an analytical Aurora Frontier™ column (60 cm × 75 μm, 1.7 μm C18, AUR3-60075C18-CSI, Ion Opticks) with a flow rate of 0.3 μL/min for a total duration of 130 min and ionized at 1.8 kV in the positive ion mode. The gradient was composed of 6% solvent B (7 min), 6-25% B (88 min), 25-40% B (25 min), 40–98% B (1 min) and 98% B (9min); solvent A: 2% acetonitrile (CAN) and 0.2% formic acid (FA) in water; solvent B: 80% ACN and 0.2% FA. MS1 scans were acquired at the resolution of 60,000 from 375 to 1500 m/z, AGC target 3e6, and maximum injection time 15 ms. The 12 most abundant ions in MS2 scans were acquired at a resolution of 30,000, AGC target 1e5, maximum injection time 45 ms, and normalized collision energy of 28. Dynamic exclusion was set to 30 s and ions with charge +1, +7, +8 and >+8 were excluded. The temperature of ion transfer tube was 275°C and the S-lens RF level was set to 60.

Data was processed in Proteome Discoverer with SEQUEST (version 2.5, Thermo Scientific) against the UniProt mouse proteome (2020 version, 55,466 entries). Trypsin/P was set as the digestion enzyme, allowing a maximum of two missed cleavages. Dynamic modifications were set to oxidation on methionine (M, +15.995 Da) and protein N-terminal acetylation (+42.011 Da). Carbamidomethylation on cysteine (C, +57.021 Da) was set as a fixed modification. The maximum parental mass error was set to 10 ppm, and the MS2 mass tolerance was set to 0.05 Da. The false discovery threshold was set strictly to 0.01 using the Percolator Node validated by q-value. The relative abundance of parental peptides was calculated by integration of the area under the curve of the MS1 peaks using the Minora LFQ node. The mass tolerance used to align features across runs was set to 0.01 ppm in the Feature Mapper node. Differential expression analysis was performed using the tidyproteomics package v1.8.2.^27^ Protein abundance values were median-normalized within each sample group. Missing data was subsequently imputed using the corresponding intra-group median values.

Statistically significant differences between groups were calculated using two-sided Student t-test. The mass spectrometry proteomics data have been deposited to the ProteomeXchange Consortium via the PRIDE partner repository with the dataset identifier PXD084923.^28^ Proteins were classified broadly into several catalogs according to the Gene Ontology (GO) annotation (geneontology.org). Overrepresentation analyses of GO terms, including biological process, molecular function, and cellular component, were performed using the Enrichr database system (https://maayanlab.cloud/Enrichr/enrich), which is a molecular functional interaction database. All proteins detected in the mass-spectrometry experiments were used as background for comparison. Terms with a Benjamini–Hochberg-adjusted q value below 0.01 were considered significant.

### Statistics

Comparisons between two groups were made using a two-tailed, unpaired Student’s t test. Comparisons between several groups were made using either a one or two-way ANOVA with a multiple comparison correction as indicated for each experiment. All analyses were performed with GraphPad Prism 9 (GraphPad Software). Results were considered statistically significant if the p value was less than 0.05. IC₅₀ values were determined by nonlinear regression as described in the legend to Figure 4, and differential protein abundance was assessed using tidyproteomics described above. Experiments presented as a single time course were analyzed qualitatively, and no inferential statistical comparisons were performed.

## DATA AVAILABILITY STATEMENT

The mass spectrometry proteomics data generated in this study have been deposited to the ProteomeXchange Consortium via the PRIDE partner repository under dataset identifier PXD084923. For all other data supporting the findings of this study, qualified researchers may request access from the corresponding author by contacting.

## ACKNOWLEDGMENTS

This project was supported by the Washington University Institute of Clinical and Translational Sciences which is, in part, supported by the NIH/National Center for Advancing Translational Sciences (NCATS), CTSA grant #UL1TR002345 (ARF).

Research reported in this publication was supported by the National Institute Of Arthritis And Musculoskeletal And Skin Diseases of the National Institutes of Health under Award Numbers K08AR075894 (ARF) and R03AR081395 (ARF), and by the National Institute Of Neurological Disorders And Stroke of the National Institutes of Health under Award Number R01NS134506 (ARF, TFC, SQH). The content is solely the responsibility of the authors and does not necessarily represent the official views of the National Institutes of Health.

This project was additionally supported by the Washington University Hope Center for Neurological Disorders (ARF) The Doris Duke Fund to Retain Clinician Scientists (DDFRCS) (ARF), The LGMD-1D DNAJB6 Foundation, The Eleven Eleven Foundation for LGMDD1 Research Funding (ARF), The RTW Charitable Foundation (ARF), The Children’s Discovery Institute of Washington University and St. Louis Children’s Hospital (ARF) (Faculty Scholar Award - MI-FR-2022-1004) and the American Society of Gene and Cell therapy (ASGCT) (ARF).

Mouse behavioral work in the Animal Behavioral Core was supported by funds provided by the McDonnell Center for Systems Neuroscience, McDonnell Center for Cellular and Molecular Neuroscience, and the Taylor Family Institute at Washington University in St. Louis. The authors thank Amanda R. Titus for providing technical support through the Animal Behavioral Core.

Imaging experiments were performed in part through the use of Washington University Center for Cellular Imaging (WUCCI) and supported by Washington University School of Medicine.

## AUTHOR CONTRIBUTIONS

AV, MH, MMP, ABLD, JAD, RB, SKP, TYW, CMY, TFC: investigation, methodology, writing – review and editing. TMM, HLT, SQH: Conceptualization, writing – review and editing. CCW: Conceptualization, funding acquisition, resources, supervision, writing – review and editing. ARF: Conceptualization, formal analysis, funding acquisition, investigation, methodology, resources, supervision, validation, visualization, writing original draft, writing – review and editing.

## DECLARATION OF GENERATIVE AI AND AI-ASSISTED TECHNOLOGIES IN THE WRITING PROCESS

During the preparation of this work the authors used Claude in order to improve readability and language. After using this tool, the authors reviewed and edited the content as needed and take full responsibility for the content of the publication.

---

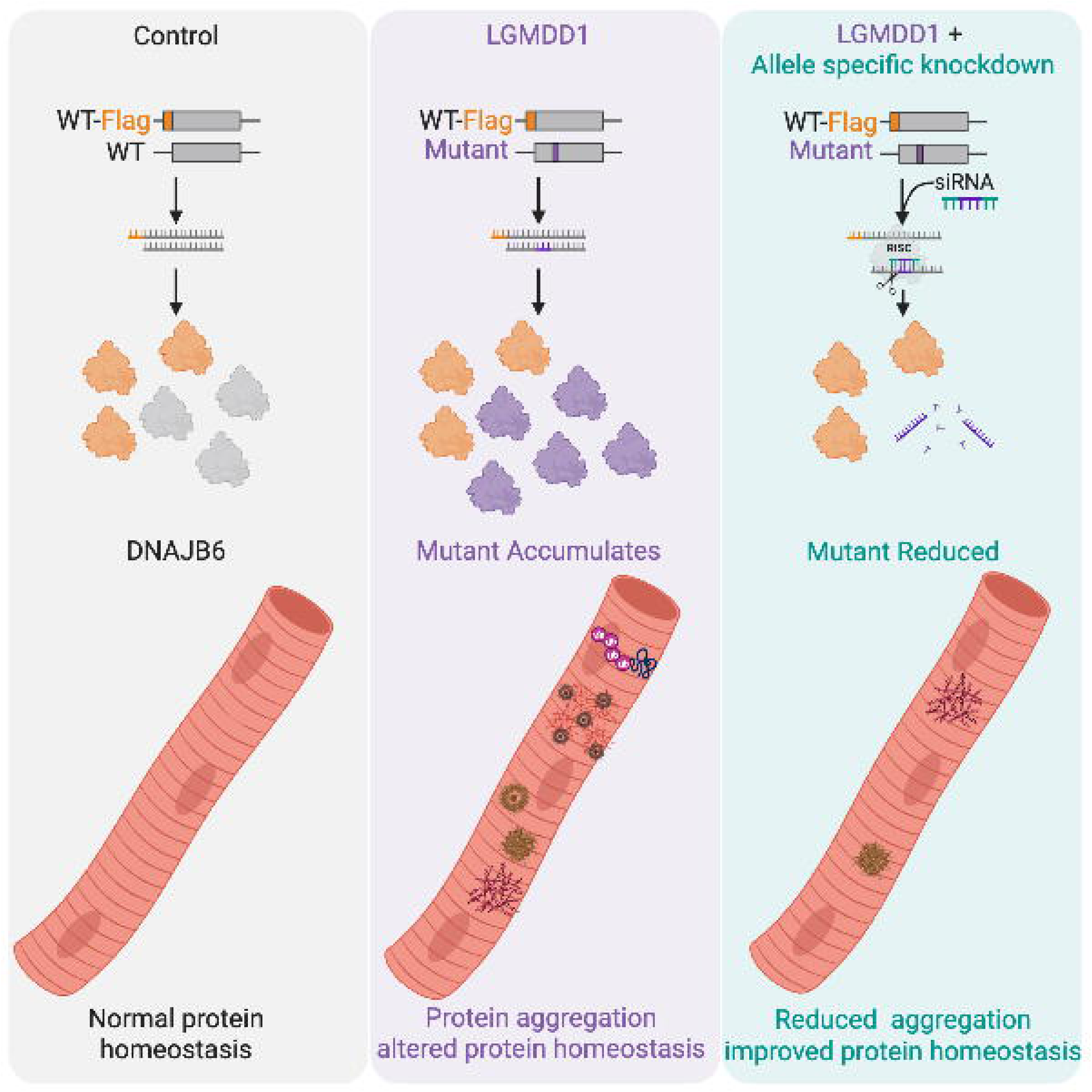

