## Supplementary material for "Allele-selective knockdown of mutant DNAJB6 improves molecular features in a primary mouse myotube model of LGMDD1": Table S1, Table S2, Figure S1

### Supplemental Tables

**Table S1.** Predicted off-target sites for first-generation siRNAs of differing length. Each candidate siRNA was tiled across the F90I variant and queried against the mouse genome (GRCm39, plus strand only) and the RefSeq mouse RNA database (2025, plus strand only). Values are the number of sites containing the indicated number of mismatches. The target sequence of each siRNA is underlined within the surrounding *Dnajb6* sequence, and the nucleotide altered by the F90I variant is shown in red. Wild-type and F90I reference sequences are given above each sub-table. Counts include all predicted binding sites, including the wild-type *Dnajb6* allele. Because first-generation siRNAs are fully complementary to F90I and carry a single mismatch relative to wild-type, the wild-type allele is counted within the 1-mismatch column. –, not determined. Sites with zero mismatches were not tabulated for the 19-nucleotide series. Totals and RefSeq RNA counts were not determined for the 19-nucleotide series.

#### (A) 22-nucleotide siRNAs

```
WT      TGGAATTCATTTTGACAGTCCATTTGAGTTTGGCTTCACATTCCGGAACCCAGATGA
F90I    TGGAATTCATTTTGACAGTCCATTTGAGATTGGCTTCACATTCCGGAACCCAGATGA
```

| siRNA | Target sequence (5'→3') | Mouse genome (GRCm39) | | | | | RefSeq mouse RNA (2025) | | | |
|---|---|---|---|---|---|---|---|---|---|---|
| | | 0 | 1 | 2 | 3 | Total | 0 | 1 | 2 | 3 |
| 1 | TGGAATTCATTTTGACAGTCCATTTGAGATTGGCTTCACATTCCGGAACCCAGATGA | 0 | 2 | 0 | 14 | 16 | 0 | 1 | 0 | 1 |
| 2 | TGGAATTCATTTTGACAGTCCATTTGAGATTGGCTTCACATTCCGGAACCCAGATGA | 0 | 2 | 0 | 16 | 18 | 0 | 1 | 0 | 0 |
| 3 | TGGAATTCATTTTGACAGTCCATTTGAGATTGGCTTCACATTCCGGAACCCAGATGA | 0 | 2 | 0 | 22 | 24 | 0 | 1 | 0 | 1 |
| 4 | TGGAATTCATTTTGACAGTCCATTTGAGATTGGCTTCACATTCCGGAACCCAGATGA | 0 | 2 | 1 | 16 | 19 | 0 | 1 | 0 | 2 |
| 5 | TGGAATTCATTTTGACAGTCCATTTGAGATTGGCTTCACATTCCGGAACCCAGATGA | 0 | 2 | 1 | 19 | 22 | 0 | 1 | 0 | 0 |
| 6 | TGGAATTCATTTTGACAGTCCATTTGAGATTGGCTTCACATTCCGGAACCCAGATGA | 0 | 2 | 1 | 20 | 23 | 0 | 1 | 0 | 1 |
| 7 | TGGAATTCATTTTGACAGTCCATTTGAGATTGGCTTCACATTCCGGAACCCAGATGA | 0 | 2 | 3 | 124 | 129 | 0 | 1 | 0 | 1 |
| 8 | TGGAATTCATTTTGACAGTCCATTTGAGATTGGCTTCACATTCCGGAACCCAGATGA | 0 | 2 | 4 | 100 | 106 | 0 | 1 | 0 | 1 |
| 9 | TGGAATTCATTTTGACAGTCCATTTGAGATTGGCTTCACATTCCGGAACCCAGATGA | 0 | 2 | 5 | 174 | 181 | 0 | 1 | 0 | 3 |
| 10 | TGGAATTCATTTTGACAGTCCATTTGAGATTGGCTTCACATTCCGGAACCCAGATGA | 0 | 2 | 4 | 126 | 132 | 0 | 1 | 0 | 0 |
| 11 | TGGAATTCATTTTGACAGTCCATTTGAGATTGGCTTCACATTCCGGAACCCAGATGA | 0 | 2 | 3 | 101 | 106 | 0 | 1 | 0 | 2 |
| 12 | TGGAATTCATTTTGACAGTCCATTTGAGATTGGCTTCACATTCCGGAACCCAGATGA | 0 | 3 | 2 | 67 | 72 | 0 | 1 | 0 | 2 |
| 13 | TGGAATTCATTTTGACAGTCCATTTGAGATTGGCTTCACATTCCGGAACCCAGATGA | 0 | 2 | 2 | 121 | 125 | 0 | 1 | 0 | 4 |
| 14 | TGGAATTCATTTTGACAGTCCATTTGAGATTGGCTTCACATTCCGGAACCCAGATGA | 0 | 2 | 3 | 146 | 151 | 0 | 1 | 0 | 2 |
| 15 | TGGAATTCATTTTGACAGTCCATTTGAGATTGGCTTCACATTCCGGAACCCAGATGA | 0 | 2 | 2 | 87 | 91 | 0 | 1 | 0 | 4 |
| 16 | TGGAATTCATTTTGACAGTCCATTTGAGATTGGCTTCACATTCCGGAACCCAGATGA | 0 | 2 | 3 | 57 | 62 | 0 | 1 | 0 | 2 |
| 17 | TGGAATTCATTTTGACAGTCCATTTGAGATTGGCTTCACATTCCGGAACCCAGATGA | 0 | 2 | 3 | 45 | 50 | 0 | 1 | 0 | 2 |
| 18 | TGGAATTCATTTTGACAGTCCATTTGAGATTGGCTTCACATTCCGGAACCCAGATGA | 0 | 3 | 1 | 65 | 69 | 0 | 1 | 0 | 0 |
| 19 | TGGAATTCATTTTGACAGTCCATTTGAGATTGGCTTCACATTCCGGAACCCAGATGA | 0 | 3 | 5 | 144 | 152 | 0 | 1 | 0 | 3 |
| 20 | TGGAATTCATTTTGACAGTCCATTTGAGATTGGCTTCACATTCCGGAACCCAGATGA | 0 | 3 | 1 | 190 | 194 | 0 | 1 | 0 | 4 |
| 21 | TGGAATTCATTTTGACAGTCCATTTGAGATTGGCTTCACATTCCGGAACCCAGATGA | 0 | 3 | 1 | 114 | 118 | 0 | 1 | 0 | 1 |
| 22 | TGGAATTCATTTTGACAGTCCATTTGAGATTGGCTTCACATTCCGGAACCCAGATGA | 0 | 3 | 2 | 105 | 110 | 0 | 1 | 1 | 1 |

#### (B) 21-nucleotide siRNAs

| siRNA | Target sequence (5'→3') | Mouse genome (GRCm39) | | | | | RefSeq mouse RNA (2025) | | | |
|---|---|---|---|---|---|---|---|---|---|---|
| | | 0 | 1 | 2 | 3 | Total | 0 | 1 | 2 | 3 |
| 1 | TGGAATTCATTTTGACAGTCCATTTGAGATTGGCTTCACATTCCGGAACCCAGATGA | 0 | 2 | 1 | 46 | 49 | 0 | 1 | 0 | 2 |
| 2 | TGGAATTCATTTTGACAGTCCATTTGAGATTGGCTTCACATTCCGGAACCCAGATGA | 0 | 2 | 0 | 60 | 62 | 0 | 1 | 0 | 2 |

| siRNA | Target sequence (5'→3') | Mouse genome (GRCm39) | | | | | RefSeq mouse RNA (2025) | | | |
|---|---|---|---|---|---|---|---|---|---|---|
| | | 0 | 1 | 2 | 3 | Total | 0 | 1 | 2 | 3 |
| 3 | TGGAATTCATTTTGACAGTCCATTTGAGATTGGCTTCACATTCCGGAACCCAGATGA | 0 | 2 | 2 | 78 | 82 | 0 | 1 | 0 | 5 |
| 4 | TGGAATTCATTTTGACAGTCCATTTGAGATTGGCTTCACATTCCGGAACCCAGATGA | 0 | 2 | 3 | 41 | 46 | 0 | 1 | 0 | 2 |
| 5 | TGGAATTCATTTTGACAGTCCATTTGAGATTGGCTTCACATTCCGGAACCCAGATGA | 0 | 2 | 4 | 71 | 77 | 0 | 1 | 0 | 4 |
| 6 | TGGAATTCATTTTGACAGTCCATTTGAGATTGGCTTCACATTCCGGAACCCAGATGA | 0 | 2 | 7 | 263 | 272 | 0 | 1 | 0 | 3 |
| 7 | TGGAATTCATTTTGACAGTCCATTTGAGATTGGCTTCACATTCCGGAACCCAGATGA | 0 | 2 | 8 | 412 | 422 | 0 | 1 | 0 | 7 |
| 8 | TGGAATTCATTTTGACAGTCCATTTGAGATTGGCTTCACATTCCGGAACCCAGATGA | 0 | 2 | 16 | 453 | 471 | 0 | 1 | 0 | 5 |
| 9 | TGGAATTCATTTTGACAGTCCATTTGAGATTGGCTTCACATTCCGGAACCCAGATGA | 0 | 2 | 19 | 405 | 426 | 0 | 1 | 0 | 7 |
| 10 | TGGAATTCATTTTGACAGTCCATTTGAGATTGGCTTCACATTCCGGAACCCAGATGA | 0 | 2 | 10 | 264 | 276 | 0 | 1 | 0 | 8 |
| 11 | TGGAATTCATTTTGACAGTCCATTTGAGATTGGCTTCACATTCCGGAACCCAGATGA | 0 | 4 | 8 | 260 | 272 | 0 | 1 | 0 | 8 |
| 12 | TGGAATTCATTTTGACAGTCCATTTGAGATTGGCTTCACATTCCGGAACCCAGATGA | 0 | 3 | 7 | 266 | 276 | 0 | 1 | 0 | 8 |
| 13 | TGGAATTCATTTTGACAGTCCATTTGAGATTGGCTTCACATTCCGGAACCCAGATGA | 0 | 2 | 9 | 323 | 334 | 0 | 1 | 0 | 7 |
| 14 | TGGAATTCATTTTGACAGTCCATTTGAGATTGGCTTCACATTCCGGAACCCAGATGA | 0 | 2 | 11 | 389 | 402 | 0 | 1 | 0 | 11 |
| 15 | TGGAATTCATTTTGACAGTCCATTTGAGATTGGCTTCACATTCCGGAACCCAGATGA | 0 | 2 | 6 | 258 | 266 | 0 | 1 | 0 | 9 |
| 16 | TGGAATTCATTTTGACAGTCCATTTGAGATTGGCTTCACATTCCGGAACCCAGATGA | 0 | 2 | 5 | 125 | 132 | 0 | 1 | 0 | 2 |
| 17 | TGGAATTCATTTTGACAGTCCATTTGAGATTGGCTTCACATTCCGGAACCCAGATGA | 0 | 3 | 3 | 242 | 248 | 0 | 1 | 0 | 4 |
| 18 | TGGAATTCATTTTGACAGTCCATTTGAGATTGGCTTCACATTCCGGAACCCAGATGA | 0 | 3 | 16 | 309 | 328 | 0 | 1 | 0 | 7 |
| 19 | TGGAATTCATTTTGACAGTCCATTTGAGATTGGCTTCACATTCCGGAACCCAGATGA | 0 | 3 | 20 | 604 | 627 | 0 | 1 | 0 | 8 |
| 20 | TGGAATTCATTTTGACAGTCCATTTGAGATTGGCTTCACATTCCGGAACCCAGATGA | 0 | 3 | 4 | 401 | 408 | 0 | 1 | 0 | 8 |
| 21 | TGGAATTCATTTTGACAGTCCATTTGAGATTGGCTTCACATTCCGGAACCCAGATGA | 0 | 3 | 10 | 370 | 383 | 0 | 1 | 0 | 7 |

#### (C) 19-nucleotide siRNAs

| siRNA | Target sequence (5'→3') | Mouse genome (GRCm39) | | |
|---|---|---|---|---|
| | | 0 | 1 | 2 |
| 1 | TGGAATTCATTTTGACAGTCCATTTGAGATTGGCTTCACATTCCGGAACCCAGATGA | 0 | 2 | 11 |
| 2 | TGGAATTCATTTTGACAGTCCATTTGAGATTGGCTTCACATTCCGGAACCCAGATGA | – | 2 | 18 |
| 3 | TGGAATTCATTTTGACAGTCCATTTGAGATTGGCTTCACATTCCGGAACCCAGATGA | – | 3 | 26 |
| 4 | TGGAATTCATTTTGACAGTCCATTTGAGATTGGCTTCACATTCCGGAACCCAGATGA | – | 6 | 71 |
| 5 | TGGAATTCATTTTGACAGTCCATTTGAGATTGGCTTCACATTCCGGAACCCAGATGA | – | 5 | 103 |
| 6 | TGGAATTCATTTTGACAGTCCATTTGAGATTGGCTTCACATTCCGGAACCCAGATGA | – | 3 | 168 |
| 7 | TGGAATTCATTTTGACAGTCCATTTGAGATTGGCTTCACATTCCGGAACCCAGATGA | – | 5 | 189 |
| 8 | TGGAATTCATTTTGACAGTCCATTTGAGATTGGCTTCACATTCCGGAACCCAGATGA | – | 4 | 131 |
| 9 | TGGAATTCATTTTGACAGTCCATTTGAGATTGGCTTCACATTCCGGAACCCAGATGA | – | 7 | 97 |
| 10 | TGGAATTCATTTTGACAGTCCATTTGAGATTGGCTTCACATTCCGGAACCCAGATGA | – | 5 | 132 |
| 11 | TGGAATTCATTTTGACAGTCCATTTGAGATTGGCTTCACATTCCGGAACCCAGATGA | – | 4 | 108 |
| 12 | TGGAATTCATTTTGACAGTCCATTTGAGATTGGCTTCACATTCCGGAACCCAGATGA | – | 3 | 100 |
| 13 | TGGAATTCATTTTGACAGTCCATTTGAGATTGGCTTCACATTCCGGAACCCAGATGA | – | 3 | 104 |
| 14 | TGGAATTCATTTTGACAGTCCATTTGAGATTGGCTTCACATTCCGGAACCCAGATGA | – | 3 | 67 |
| 15 | TGGAATTCATTTTGACAGTCCATTTGAGATTGGCTTCACATTCCGGAACCCAGATGA | – | 3 | 68 |
| 16 | TGGAATTCATTTTGACAGTCCATTTGAGATTGGCTTCACATTCCGGAACCCAGATGA | – | 5 | 68 |
| 17 | TGGAATTCATTTTGACAGTCCATTTGAGATTGGCTTCACATTCCGGAACCCAGATGA | – | 4 | 138 |
| 18 | TGGAATTCATTTTGACAGTCCATTTGAGATTGGCTTCACATTCCGGAACCCAGATGA | – | 4 | 121 |
| 19 | TGGAATTCATTTTGACAGTCCATTTGAGATTGGCTTCACATTCCGGAACCCAGATGA | – | 5 | 105 |

**Table S2.** Predicted binding sites for second-generation siRNAs. Second-generation siRNAs were derived from siRNA 11 by introducing one additional mismatch at successive positions across the sequence; no siRNA 11-11 exists because position 11 already carries the first-generation mismatch. Sequences were queried against the RefSeq mouse RNA database (2025) for transcripts containing the indicated number of mismatches, and by NCBI BLAST for transcripts containing at least 15 contiguous complementary nucleotides. Counts include all predicted binding sites, including the wild-type *Dnajb6* transcript; because second-generation siRNAs carry two mismatches relative to wild-type, the wild-type transcript is counted within the 2-mismatch column. Genes listed in the final column are those containing a site with at least 15 contiguous complementary nucleotides. Nucleotides shown in red differ from the wild-type sequence. Bold sequences indicate the six candidates selected for dose-response testing. n.a., not analyzed. Binding-site analysis was performed only for the six candidate siRNAs advanced to dose-response testing; remaining sequences are listed for completeness.

| Oligo name | Sequence (5'→3') | 0 | 1 | 2 | 3 | ≥15 contiguous bp | Off-target gene(s) |
|---|---|---|---|---|---|---|---|
| mDNAJB6_22_F90I_11-1 | ACCATTTGAGATTGGCTTCACA | n.a. | n.a. | n.a. | n.a. | n.a. | n.a. |
| mDNAJB6_22_F90I_11-2 | **TGCATTTGAGATTGGCTTCACA** | 0 | 0 | 1 | 0 | 1 | ATP2c2 |
| mDNAJB6_22_F90I_11-3 | **TCGATTTGAGATTGGCTTCACA** | 0 | 0 | 1 | 0 | 2 | ATP2C2, ARHGAP20 |
| mDNAJB6_22_F90I_11-4 | **TCCTTTTGAGATTGGCTTCACA** | 0 | 0 | 1 | 2 | 0 | none |
| mDNAJB6_22_F90I_11-5 | **TCCAATTGAGATTGGCTTCACA** | 0 | 0 | 1 | 0 | 1 | TAS2R113 |
| mDNAJB6_22_F90I_11-6 | **TCCATATGAGATTGGCTTCACA** | 0 | 0 | 1 | 3 | 1 | PLA2G4A |
| mDNAJB6_22_F90I_11-7 | TCCATTAGAGATTGGCTTCACA | n.a. | n.a. | n.a. | n.a. | n.a. | n.a. |
| mDNAJB6_22_F90I_11-8 | TCCATTTCAGATTGGCTTCACA | n.a. | n.a. | n.a. | n.a. | n.a. | n.a. |
| mDNAJB6_22_F90I_11-9 | TCCATTTGTGATTGGCTTCACA | n.a. | n.a. | n.a. | n.a. | n.a. | n.a. |
| mDNAJB6_22_F90I_11-10 | TCCATTTGACATTGGCTTCACA | n.a. | n.a. | n.a. | n.a. | n.a. | n.a. |
| mDNAJB6_22_F90I_11-12 | TCCATTTGAGAATGGCTTCACA | n.a. | n.a. | n.a. | n.a. | n.a. | n.a. |
| mDNAJB6_22_F90I_11-13 | **TCCATTTGAGATAGGCTTCACA** | 0 | 0 | 1 | 0 | 0 | none |
| mDNAJB6_22_F90I_11-14 | TCCATTTGAGATTCGCTTCACA | n.a. | n.a. | n.a. | n.a. | n.a. | n.a. |
| mDNAJB6_22_F90I_11-15 | TCCATTTGAGATTGCCTTCACA | n.a. | n.a. | n.a. | n.a. | n.a. | n.a. |
| mDNAJB6_22_F90I_11-16 | TCCATTTGAGATTGGGTTCACA | n.a. | n.a. | n.a. | n.a. | n.a. | n.a. |
| mDNAJB6_22_F90I_11-17 | TCCATTTGAGATTGGCATCACA | n.a. | n.a. | n.a. | n.a. | n.a. | n.a. |
| mDNAJB6_22_F90I_11-18 | TCCATTTGAGATTGGCTACACA | n.a. | n.a. | n.a. | n.a. | n.a. | n.a. |
| mDNAJB6_22_F90I_11-19 | TCCATTTGAGATTGGCTTGACA | n.a. | n.a. | n.a. | n.a. | n.a. | n.a. |
| mDNAJB6_22_F90I_11-20 | TCCATTTGAGATTGGCTTCTCA | n.a. | n.a. | n.a. | n.a. | n.a. | n.a. |
| mDNAJB6_22_F90I_11-21 | TCCATTTGAGATTGGCTTCAGA | n.a. | n.a. | n.a. | n.a. | n.a. | n.a. |
| mDNAJB6_22_F90I_11-22 | TCCATTTGAGATTGGCTTCACT | n.a. | n.a. | n.a. | n.a. | n.a. | n.a. |

**Table S3.** Proteomic analysis of F90I and ΔGF DNAJB6 myotubes. Provided as a separate Microsoft Excel file. $Log_2$ fold changes, p values, and adjusted p values for all proteins quantified in primary myotubes from *Dnajb6*$^{WT/F90I}$ versus *Dnajb6*$^{WT/WT}$ mice, and in *Dnajb6*$^{WT/WT}$ myotubes treated with the ΔGF antisense oligonucleotide versus control oligonucleotide.

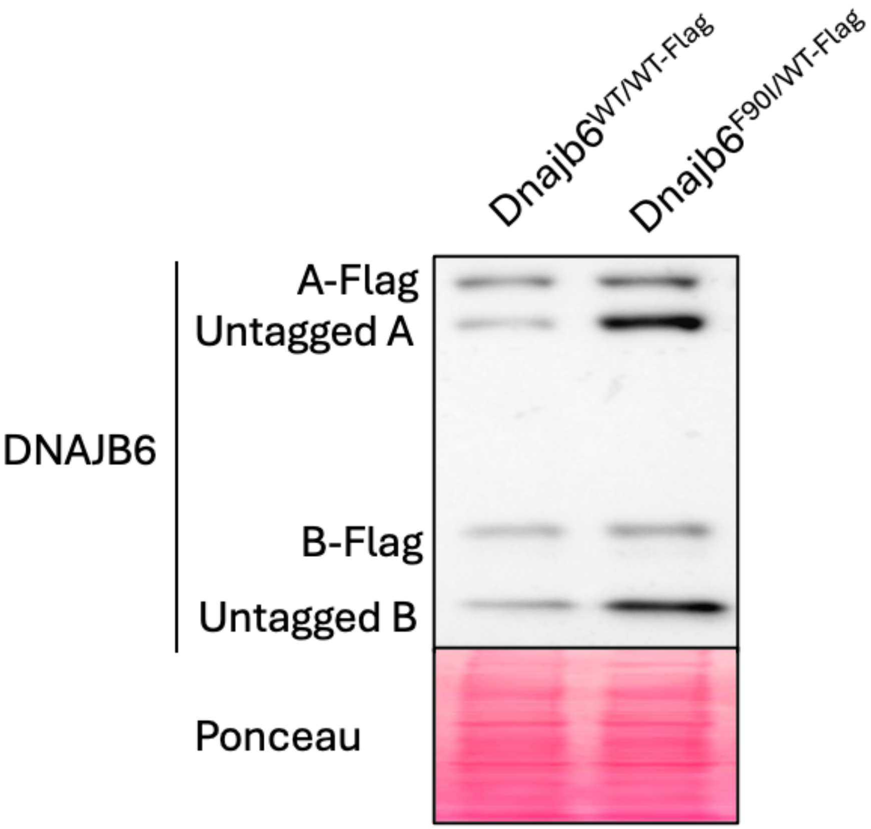


**Figure S1.** FLAG tagging does not substantially alter DNAJB6 abundance. Anti-DNAJB6 immunoblot of primary myotubes from *Dnajb6*$^{WT/WT\text{-}Flag}$ and *Dnajb6*$^{F90I/WT\text{-}Flag}$ mice differentiated for 3 days (see Methods). FLAG-tagged DNAJB6a and DNAJB6b (A-Flag, B-Flag) migrate above their untagged counterparts. Ponceau S staining was used to assess protein loading. A single experiment is shown; densitometry was not performed and no inferential statistical comparisons were made.
